# When should adaptation arise from a polygenic response versus few large effect changes?

**DOI:** 10.1101/2025.05.15.654234

**Authors:** William R Milligan, Laura K Hayward, Guy Sella

**Affiliations:** Department of Biological Sciences, Columbia University, New York, NY, USA; Institute of Science and Technology Austria, Klosterneuburg, Austria; Program for Mathematical Genomics, Columbia University, New York, NY, USA

## Abstract

The question of when adaptation involves genetic changes of large effect versus a polygenic response traces back to the foundation of evolutionary biology. While there are compelling reasons to expect polygenic adaptation to be common, direct evidence is still lacking. In turn, there are hundreds of examples of large effect adaptations across species, but it is unclear whether they are a common occurrence in any given species. Synthesizing the different lines of evidence is further complicated by differences in study designs. Here, we reframe this long-standing question in terms of properties of the trait under selection. We ask how the genetic basis of adaptation is expected to depend on key aspects of the genetic variation in the trait and on the extent of changes in selection pressures on it (i.e., the “trait’s ecology”). We consider a quantitative trait subject to stabilizing selection and model the response to selection when a population at mutation-selection-drift balance experiences a sudden shift in the optimal value. Using this model, we delimit how the contributions of large effect and polygenic changes to adaptation depend on the genetics and ecology of the trait, as well as on other salient factors. We thus formulate testable predictions about when different modes of adaptation are expected and outline a framework within which to interpret disparate sources of evidence about the genetic basis of adaptation.

## Introduction

Questions about the workings of adaptation, notably, whether it typically arises from numerous changes with tiny effects or from few changes with large ones, trace back to early debates surrounding Darwin’s “Origin of Species” and the bitter conflict between early geneticists and biometricians (Darwin 1859; Huxley 1860; Orr and Coyne 1992; Provine 2001). Geneticists argued that evolution proceeds via mutations with large phenotypic effects (“saltations”) of the kind observed in Mendel’s experiments (Bateson 1913; Morgan 1932). Biometricians instead emphasized gradual, nearly continuous phenotypic change that aligned with the apparent continuous heritable variation in quantitative traits (i.e., traits that take continuous values) and was observed in the response to artificial selection on these traits (Pearson 1898). While these views of inheritance were reconciled during the Modern Synthesis (Fisher 1918), the debate about adaptation was never resolved. Nonetheless, for nearly a century, quantitative geneticists have assumed that heritable variation in traits is highly polygenic, and that adaptative changes in these traits take a continuous, highly polygenic form (Falconer and McKay 1996; Walsh and Lynch 2018). In population genetics, this view has been more contentious. Notably, Orr and Coyne (1992) argued that the handful of clear cases of adaptation known at the time involved large effect genetic changes, whereas the case for polygenic adaptation relied on debatable theoretical arguments and indirect and inconclusive evidence.

We have learned a lot since Orr and Coyne’s review. The degree of polygenicity of heritable variation in many traits has become evident recently, at least in humans, where this question has been studied systemically in huge samples. Evidence from genome-wide association studies (GWAS) indicates that heritable variation in many traits of different kinds is widely distributed in the genome and thinly spread over numerous variants (Yang et al. 2010; Loh et al. 2015; Shi et al. 2016; Sinnott-Armstrong et al. 2021; Yengo et al. 2022). Estimates of the number of causal variants affecting traits range from a few thousands for “simple” traits, such as the biomarkers measured in a standard blood panel, to more than 100,000 for traits such as height and body mass index (Yang et al. 2010; Zhang et al. 2018; Frei et al. 2019; O’Connor et al. 2019; Simons et al. 2025). If traits are typically so highly polygenic, then adaptive changes in them should be accomplished through tiny changes to allele frequencies at the many variants that affect them (Fisher 1930; Wright 1931). Thus, in humans at least, we would expect polygenic adaptation to be common (Pritchard et al. 2010; Pritchard and Di Rienzo 2010).

Identifying the footprints of such polygenic adaptation is extremely difficult, however. As the tiny changes to frequencies of individual variants that affect a given trait could easily be obfuscated by genetic drift and pleiotropic selective effects, evidence for polygenic adaptation has been garnered by pooling many variants associated with that trait in GWAS (Turchin et al. 2012; Berg and Coop 2014; Robinson et al. 2015; Field et al. 2016; Speidel et al. 2019). This approach, however, is sensitive to even subtle biases in GWAS due to population structure confounding (Sohail et al. 2019; Barton et al. 2019; Berg et al. 2019; Smith et al. 2025).

Meanwhile, we now have hundreds of examples of large effect adaptive changes distributed over many species (Courtier-Orgogozo et al. 2020). Well studied cases include, e.g., coding changes to the sodium potassium ATPase gene underlying cardenolides resistance in the monarch caterpillar and many other species (Price and Lingrel 1988; Holzinger and Wink 1996; Mohammadi et al. 2022; Agrawal et al. 2024); the repeated loss of function of *FRIGIDA* in *Arabidopsis thaliana* and other plants in response to selection on flowering time (Le Corre et al. 2002; Stinchcombe et al. 2004; Shindo et al. 2005; Toomajian et al. 2006; Kuittinen et al. 2008); changes to EDA and PitX that lead to adaptive loss of body armor and pelvic spine, respectively, in threespine sticklebacks living in freshwater habitats (Colosimo et al. 2005; Chan et al. 2010; Jones et al. 2012); and recurrent regulatory changes in the LCT gene that confer lactase persistence in different human populations (Enattah et al. 2002; Swallow 2003; Tishkoff et al. 2007; Imtiaz et al. 2007; Ingram et al. 2009).

While examples of large effect adaptations continue to amass, their relative importance in evolution remains elusive. It is easier to identify large effect adaptive changes than it is polygenic adaptation, so the ascertainment is clearly biased toward their discovery. More generally, the approaches taken to study adaptation vary in the questions they ask, in their power to detect different modes of adaptation, and in their biases. Therefore, it remains unclear if large effect changes represent a common mode of adaptation in any given species (see, e.g., Coop et al. 2009; Pritchard et al. 2010; Pritchard and Di Rienzo 2010; Hernandez et al. 2011; Elyashiv et al. 2016; Murphy et al. 2022).

How then can we gain a broad understanding of the genetic basis of adaptation, mindful of the different approaches and limitations of studies to date? We propose that theory offers a way forward. To this end, we reframe the question in terms of the trait under selection and ask how the genetic basis of adaptation is expected to depend on key properties of the genetic variation in the trait and on the changes in selection pressures that act on it—on the genetics and ecology of the trait.

We study this question under classic, empirically supported assumptions. Motivated by the evidence that many traits are subject to stabilizing selection (Johnson and Barton 2005; Walsh and Lynch 2018; Sella and Barton 2019)—with fitness declining with displacement from an optimal trait value—we consider the adaptive response to a sudden shift in optimal value. In this setting, the magnitude of the shift reflects a key aspect of the ecology of the trait, namely how different the selection pressure of the new environment is relative to the old one. Given extensive evidence that heritable phenotypic variation in many simple and complex traits is predominated by mutation-selection-drift balance (MSDB) (Jobling et al. 2013; Walsh and Lynch 2018; Sella and Barton 2019), we assume that the population is at MSDB before the shift in optimum occurs. In this setting, selection on variants both before and after the shift derive from selection on the trait, and the genetic architecture of the trait at MSDB can be characterized in terms of the (population-scaled) rate of mutations that affect it and the distribution of their effects sizes (Keightley and Hill 1988; Simons et al. 2018). We therefore consider these parameters or equivalent ones as a description of the genetics of the trait. Thus, we ask how the dynamics and adaptive contributions of large effect and polygenic changes depend on the genetics and ecology of the trait.

Our investigation builds upon previous work that made similar assumptions. Lande (1983) studied the adaptive response to a sudden shift in trait optimum, assuming an infinite population size and that genetic variation in the trait arises from a large effect (major) allele and a normally distributed “Fisherian” (infinitesimal) genetic background. Chevin and Hospital (2008) considered a similar scenario in a finite population and focused on the dynamic of the major allele, in particular on whether it fixes and the sweep signatures it generates when it does. To this end, they assumed that the major allele’s initial frequency (before the shift in optimum) takes a particular value, which allowed them to neglect the effects on genetic drift on its trajectory. While both studies highlighted that the initial frequency of the major allele critically affects whether it would make a long-term adaptive contribution, neither considered how selection and genetic drift before the shift in optimum would affect this frequency. Nor did they consider what would happen if there were more than one major allele (Orr and Coyne 1992). Recent work by Höllinger et al. (2023) accounted for these factors, assuming that the number and frequency of segregating alleles before the shift in optimum follow from their distributions at mutation-selection-drift balance (MSDB). They assumed that all alleles have the same effect size, however, so obviously did not consider contributions of few large effect changes versus many small ones. We overcome these limitations by assuming that the population starts at MSDB with a distribution of effect sizes, as we detail next.

## Results and Discussion

### The model

Figure 1 depicts our model; for a summary of notation, see Table S1, and for detailed conditions on parameters, see supplement section 1.

**Figure 1.**
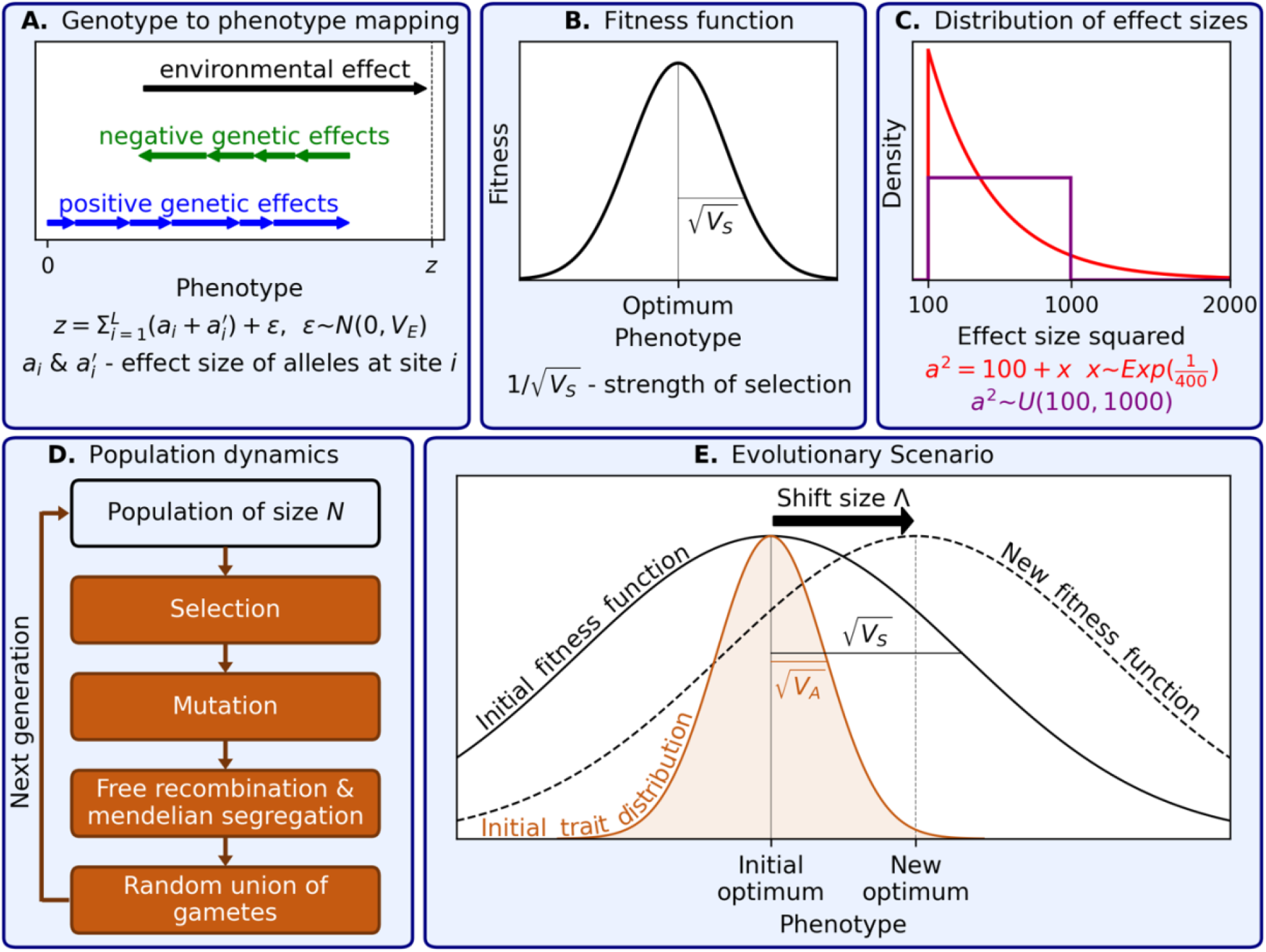
The model.

For the genotype to phenotype mapping (Figure 1A), we assume the canonical additive model from quantitative genetics: an individual’s trait value, *z*, is equal to the sum of effects of its parents’ alleles, *a* and *a*′, at the *L* loci that affect the trait, as well as a normally-distributed contribution from the environment, *ϵ*(Falconer and McKay 1996; Walsh and Lynch 2018). This model is the workhorse of quantitative genetics, and the number of loci affecting the trait, *L*, is usually assumed to be very large. Here, we will allow this number to vary, all the way from the Mendelian single locus extreme to the highly polygenic one.

Next, we model selection acting on the trait (Figure 1B). We assume that the trait is subject to stabilizing selection, namely, that fitness declines with displacement from an optimal trait value. Specifically, we assume a Gaussian fitness function, in which the reciprocal of the width, 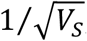, quantifies the strength of selection. So long as the population does not stray too far from the optimum, the particular shape of the fitness function does not matter (because fitness is well approximated by a quadratic form near the optimum). The environmental contribution to the trait weakens the effect of selection in a way that is equivalent to increasing the width of this Gaussian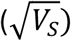 (see, e.g., Lande 1975; Turelli 1984; Bürger 2000). We can therefore consider only the genetic contribution moving forward.

When new mutations affecting the trait arise, their effects on the trait are drawn from a distribution (Figure 1C). As we detail below, the adaptive response of alleles that are effectively and nearly neutral at MSDB can be approximated in terms of their total contribution to variance in the trait, *σ*^2^. We therefore explicitly model only the mutational distribution of large effect alleles—that are strongly selected at MSDB—using one distribution here (in red) and another in Fig. S19 (in purple). We measure these effects in units of 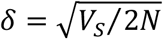, where an allele’s squared effect size in these units, *a*^2^, equals their population-scaled selection coefficient at MSDB (see below and Simons et al. 2018).

To model population genetic dynamics, we assume the standard model of a diploid, panmictic population of constant size *N*, with selection, mutation, free recombination and Mendelian segregation (Figure 1D). In simulations and analysis, we assume the infinite sites approximation, such that segregating sites are bi-allelic, and linkage equilibrium, which provides a good approximation in this model (see Hayward and Sella 2022 and supplement section 3).

Lastly, we consider a simple yet highly relevant evolutionary scenario (Figure 1E). We assume that the population begins at MSDB around the optimal phenotype, and then the optimum exhibits a sudden shift of magnitude Λ(as in Hayward and Sella 2022 and Höllinger et al. 2023). To assure that our results are insensitive to the shape of the fitness function, we assume that the shift is smaller than, or on the order of, the width of the fitness function (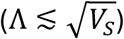); this assumption is not particularly restrictive, in that it allows for shifts of several phenotypic standard deviations unless the phenotypic variance is unrealistically large (i.e., *V*_*A*_(0) ≳ *V*_*s*_, see supplement section 1). We ask how the adaptive response depends on the trait genetics, e.g., on the genetic architecture of heritable variation before the shift, and on trait ecology, in this case, on the magnitude of the shift in optimum.

### Allele dynamic

The allele dynamic in this model is well approximated in terms of the first two moments of its frequency change in a single generation (Ewens 2004). For an allele with effect size *a* and frequency *x*, these moments are

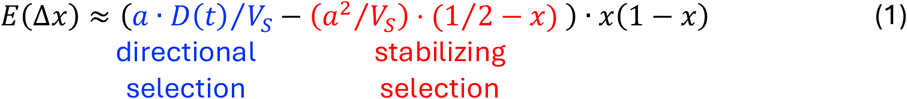

and

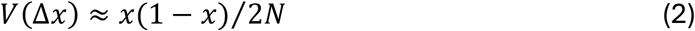

(supplement section 2 and Hayward and Sella 2022; also see Barton 1986; Burger 1991; Charlesworth 2013; de Vladar and Barton 2014; Jain and Stephan 2017). The variance in allele frequency at the bottom is the standard genetic drift term.

The expected change in allele frequency in a single generation (Eq. 1) reflects two different modes of selection on the trait: directional and stabilizing selection. Directional selection takes an additive (semi-dominant) form, with a selection coefficient that is proportional to the allele’s effect on the trait, *a*, and to the population’s mean distance from the optimum, *D*. It increases the frequency of alleles whose effects reduce the distance from the optimum and decreases the frequency of alleles with opposing effects.

While directional selection reduces the distance between the population mean and optimum, stabilizing selection reduces the phenotypic variance around the mean. Stabilizing selection takes an under-dominant form, with a selection coefficient that is proportional to the effect size squared. It reduces the frequency of the minor allele and increases the frequency of the major allele, with selection becoming weaker near frequency ½.

Before the shift in optimum, at MSDB, the population mean is at the optimum, so only the stabilizing selection term affects the allele dynamic (Simons et al. 2018). This dynamic is determined by the population-scaled selection coefficient, *s*_*e*_ = 2*Ns*_*e*_ = 2*N a*^2^⁄*V*_*s*_, which reflects the balance between selection and drift.

Akin to the more familiar semi-dominant case, the scaled selection coefficient delineates three dynamic regimes. In this case, the boundary between dynamic regimes corresponds to *s*_*e*_ ≈ 5 rather than the usual 1 (because, for a given selection coefficient, selection becomes increasingly weaker than in the semi-dominant case as the minor allele frequency increases). Effectively neutral alleles, with *s*_*e*_ ≪ 5, are dominated by drift, allowing them to ascend to high frequencies and fix. Nearly neutral alleles, with *s*_*e*_ ≈ 5, are affected to similar degrees by drift and selection, allowing them to ascend to appreciable frequencies and fix, but with lower probabilities than effectively neutral alleles. In contrast to the genetic turnover of effectively and nearly neutral alleles, the dynamic of strongly selected alleles, with *s*_*e*_ ≫ 5, is dominated by selection, which keeps them at low frequencies such that they almost never fix.

We rely on the dynamic at MSDB to define what we mean by a large effect allele. To this end, we measure the trait in units such that an allele with effect size *a* has a scaled selection coefficients *s*_*e*_ = *a*^2^ (i.e., in units of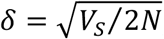). We refer to alleles with *s*_*e*_ = *a*^2^ ≪ 5 as small, to alleles with *s*_*e*_ = *a*^2^∼5 as intermediate, and to alleles with *s*_*e*_ = *a*^2^ ≫ 5 as large.

### The highly polygenic case

We first consider the adaptive response to the sudden shift in optimum in the highly polygenic case. Specifically, we assume that genetic variance in the trait is thinly spread over many segregating alleles and predominated by alleles with small and intermediate effects, and that the shift in optimum is not enormous relative to the variance in the trait before the shift (see Hayward and Sella 2022 and supplement section 4 for the exact conditions on model parameters).

The adaptive phenotypic response in this case is depicted in Figure 2A. The population mean catches up with the new optimum rapidly, because adaptation requires only minute frequency changes to the many segregating alleles that affect the trait. The genetic variance in the trait remains the approximately constant for the same reason (Figure 2B). As described by Lande (1976), the decay of the distance between the population mean and the new optimum is well approximated by

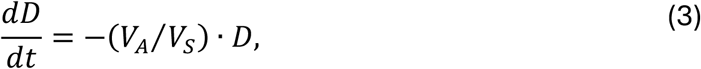

where the rate of decay is proportional to the genetic variance in the trait, *V*_*A*_, and to the strength of stabilizing selection, 1⁄*V*_*s*_. In our case, the population mean converges to the new optimum exponentially, i.e., *D*_*L*_(*t*) = Λ· *Exp*(−(*V*_*A*_⁄*V*_*s*_)*t*).

**Figure 2.**
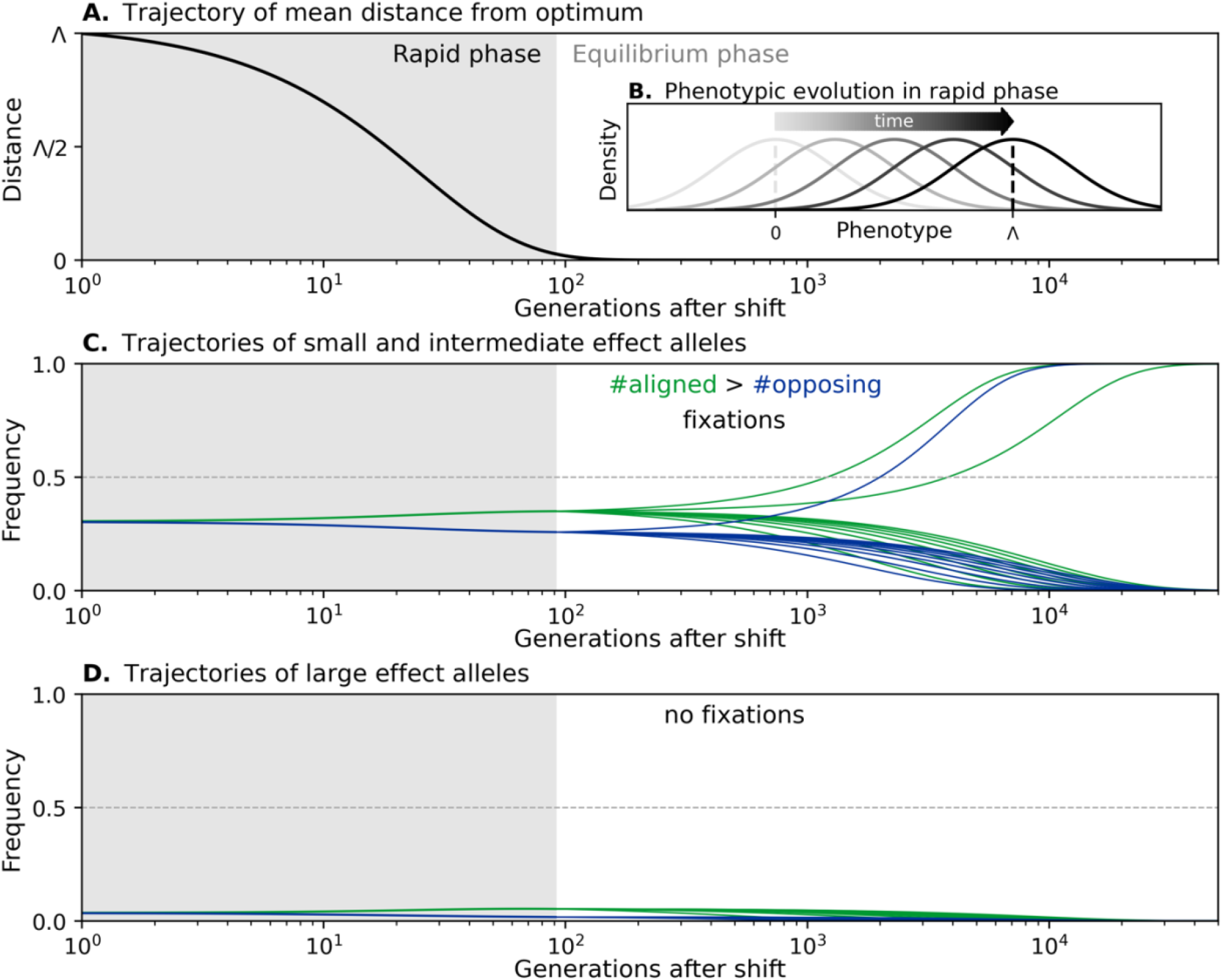
The adaptive response in the highly polygenic case. (A) The change in distance of the mean phenotype from the new optimum after the shift follows Lande’s approximation. (B) The shape of the phenotypic distribution does not change during the rapid phase. (C) The trajectories of small and intermediate effect alleles result in the preferential fixation of alleles aligned with the shift relative to those opposing it. (D) The trajectories of large effect alleles always end with their loss. These illustrations correspond to a shift size Λ= 80, variance *V*_*A*_ = 400, with cartoon trajectories corresponding to *a*^2^ = 5 and 35 and initial frequencies 1⁄*a*^2^ in (C) and (D), respectively.

Figure 2C depict the allele response that underlies phenotypic adaptation in this case, following Hayward and Sella (2022). Immediately after the shift, the mean distance from the optimum is substantial. Directional selection on the trait increases the frequency of minor alleles whose effects are aligned with the shift (in green) relative to those with opposing effects (in blue). By the end of this rapid phase, the cumulative effect of the frequency differences between aligned and opposing alleles drives the mean phenotype close to the new optimum. Because this effect is spread over many alleles, however, the frequency differences between aligned and opposing alleles remain small.

When the population mean nears the optimum the population enters an equilibration phase in which alleles are only affected by stabilizing selection and genetic drift (rather than direction selection). If the selection against minor alleles is sufficiently weak, some of them will eventually reach fixation in the long run. Since aligned minor alleles enter this equilibration phase at slightly higher frequencies than opposing alleles, more of them will fix, leading to long-term adaptation.

Importantly, large effect alleles do not fix in the population or contribute to long-term polygenic adaptation (Figure 2D). They start at low frequencies because of stabilizing selection before at shift in optimum. Directional selection during the rapid phase increases the frequency of large effect alleles whose effects are aligned with the shift relative to opposing ones. But directional selection is too short lived to push aligned alleles anywhere near frequency ½. Consequently, they are strongly selected against during the equilibration phase, and this selection pressure eventually drives all of them to extinction.

These findings, however, clarify what would have to change for large effect alleles to fix. Directional selection would have to last for long enough for them to near frequency ½. For that to happen, the shift in optimum would have to be sufficiently large and/or the genetic variance in the trait would have to be sufficiently small.

### One large effect allele and a ‘Fisherian’ background

To learn more about these conditions, we consider a case in which genetic variation in the trait has two components (akin to Lande 1983 and Chevin and Hospital 2008). One is a single segregating large effect allele, with effect size *a* and initial frequency *x*_0_ that is drawn from the frequency distribution at MSDB. The other is a normally distributed ‘Fisherian’ genetic background with variance *σ*^2^, which captures the contribution of small and intermediate effect alleles, and whose effect on adaptation follows Lande’s approximation (Eq. 3). In this case, we would like to know under what conditions the large effect allele can fix, or more precisely, the probability that it fixes.

Figure 3A illustrates the adaptive phenotypic response in this case. The distance from the optimum starts from the shift size Λ and decays with time. Initially, when the large effect allele is rare, adaptation derives almost entirely from the genetic background and follows Lande’s approximation. When the large effect allele’s frequency increases, its contribution to adaption increases and becomes substantial relative to the contribution from the background.

**Figure 3.**
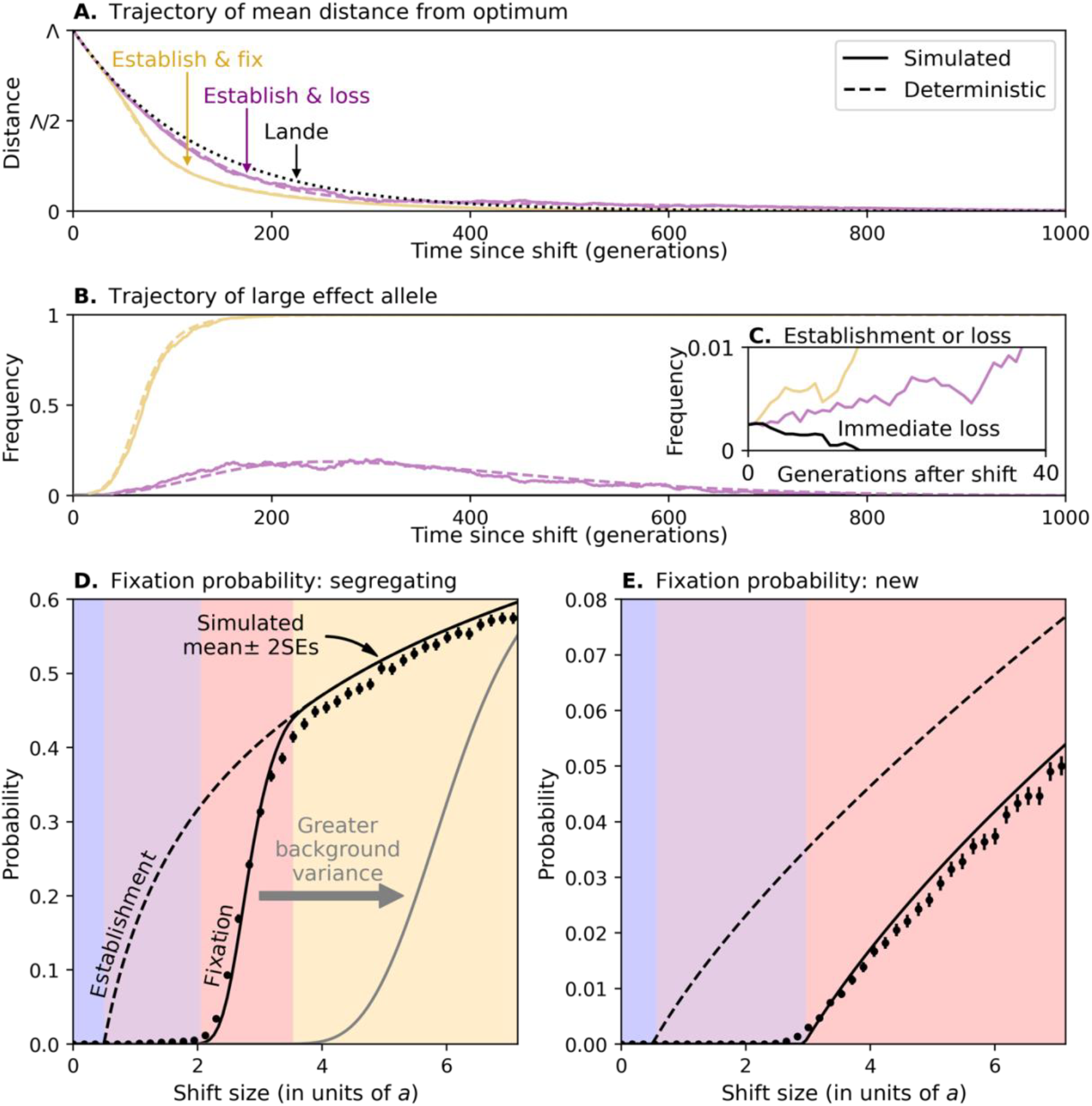
Fixation of a single large effect allele. The trajectories of the mean distance from the optimum (in A) and the trajectories of the large effect allele (in B and C) are shown for individual simulations with *N* = 5000, *a*^2^ = 200 with *x*_0_ = 1⁄(2*a*^2^), *σ*^2^ = 40, and Λ= 50 (purple) and 100 (yellow and black [in C]). The probabilities that a segregating allele establishes itself in the population and fixes (shown in D) assume *N* = 5000, *σ*^2^ = 40 (and 80 for the gray line in D), *a*^2^ = 200, and average over the distribution of initial frequencies at MSDB. The probabilities for a new allele were calculated for the same parameters, assuming that the allele is equally likely to arise (at frequency 1⁄2*N*) any time between the shift and time *t* = 606 (at which *D*_*L*_(*t*) = *δ*). These calculations are described in supplement section 5. Simulation results were averaged over 16,000 and 64,000 replicas in (D) and (E), respectively.

Figure 3B shows possible trajectories of the large effect allele. Immediately after the shift, the allele becomes beneficial. If it starts at low copy number, its initial trajectory is stochastic, and it may rapidly be lost (Figure 3C). If instead the allele ascends to some critical frequency, it is unlikely to be lost quickly. In this case, its frequency will continue to increase in a largely deterministic manner so long as directional selection is sufficiently strong. Several kinds of allelic and phenotypic trajectories can ensue (see supplement section 5.3 and Figure S5). Notably, if the allele does not make it to frequency ½ before the distance from the optimum nears 0, then when the effects of stabilizing selection on it become stronger than the effects directional selection, the selection pressure on it will reverse, and the allele will eventually be lost (Chevin and Hospital 2008). Alternatively, if the allele makes it to frequency ½ in time, then stabilizing and directional selection will likely propel it all the way to fixation. We rely on these considerations to approximate the probability that the allele fixes as the probability that it establishes itself in the population *multiplied by* the probability that it reaches frequency ½ conditional on establishing (see supplement section 5 and Figure 3D and E).

The fixation probability of a segregating allele with effect size *a* depends on the shift size (Figure 3D; also see supplement section 5.1). When shift sizes are below *a*⁄2 (in the blue region), alleles are negatively selected from the start and cannot even establish themselves in the population (Eq. 1). When shift sizes exceed *a*⁄2 (the purple region), alleles sometimes establish themselves, but they never make it to frequency ½. When shift sizes exceed some greater threshold (the red region), alleles that start at sufficiently high frequencies and establish themselves reach frequency ½ and eventually fixation. When shift sizes are even larger (the yellow region), almost all the alleles that establish themselves will fix. Increasing the contribution of the background genetic variance shortens the duration of strong directional selection, thus increasing the threshold for shift sizes that allow large effect alleles to fix in the population.

We further consider the probability of fixation of a large effect allele that arises from a mutation after the shift in optimum (Figure 3E, see supplement section 5.2). Such an allele always starts at frequency 1/2*N*, but can arise at different times. We assume that this time is uniformly distributed within the time-window in which the distance to the optimum is still substantial (specifically, while *D*_*L*_(*t*) / *δ*). Akin to the case starting from a segregating allele, when shift sizes are below *a*/2 (the blue region), the allele cannot even establish itself, whereas when shift sizes exceed some greater threshold (the red region), the allele can sometimes do so, reach frequency ½ and eventually fixation. This threshold is greater for a new allele than for a segregating one, because it always starts as a single copy, so at a lower initial frequency than a segregating allele. Moreover, even with much larger shift sizes, an allele that arises too late in the adaptive process can sometimes establish itself but can never reach fixation. This explains why the fixation probability of a new allele never converges to the probability of establishment (as it does in the yellow region in Figure 3D).

### The general case of large effect alleles and a Fisherian background

Next, we consider the general model for large effect alleles. Namely, large effect mutations arise at a rate of 2*NU* per generation, both before and after the shift in optimum, and their effect sizes are drawn from some distribution. The distribution of frequencies and effect sizes of large effect alleles that segregate before the shift derive from MSDB, whereas those that arise from mutations after the shift start from a single copy. As before, we assume a normally distributed ‘Fisherian’ background that follows Lande’s approximation, which captures the effects of small and intermediate effect alleles. This case therefore captures the range of behaviors in our model (supplement section 3.1).

Figure 4A shows how the mutational input of large effect alleles affects the fraction of the shift in optimum that their fixations account for—their long-term adaptive contribution; the rest of the shift comes from the genetic background. When the mutational input is low (in the gray shaded region corresponding to 2*NU* < 1), this contribution increases with mutational input. The reason is intuitive: increasing the number of large effect mutations increases the probability that at least one of them establishes, crosses the Rubicon at frequency ½ and eventually fixes in the population, thus contributing to long-term adaptation.

**Figure 4.**
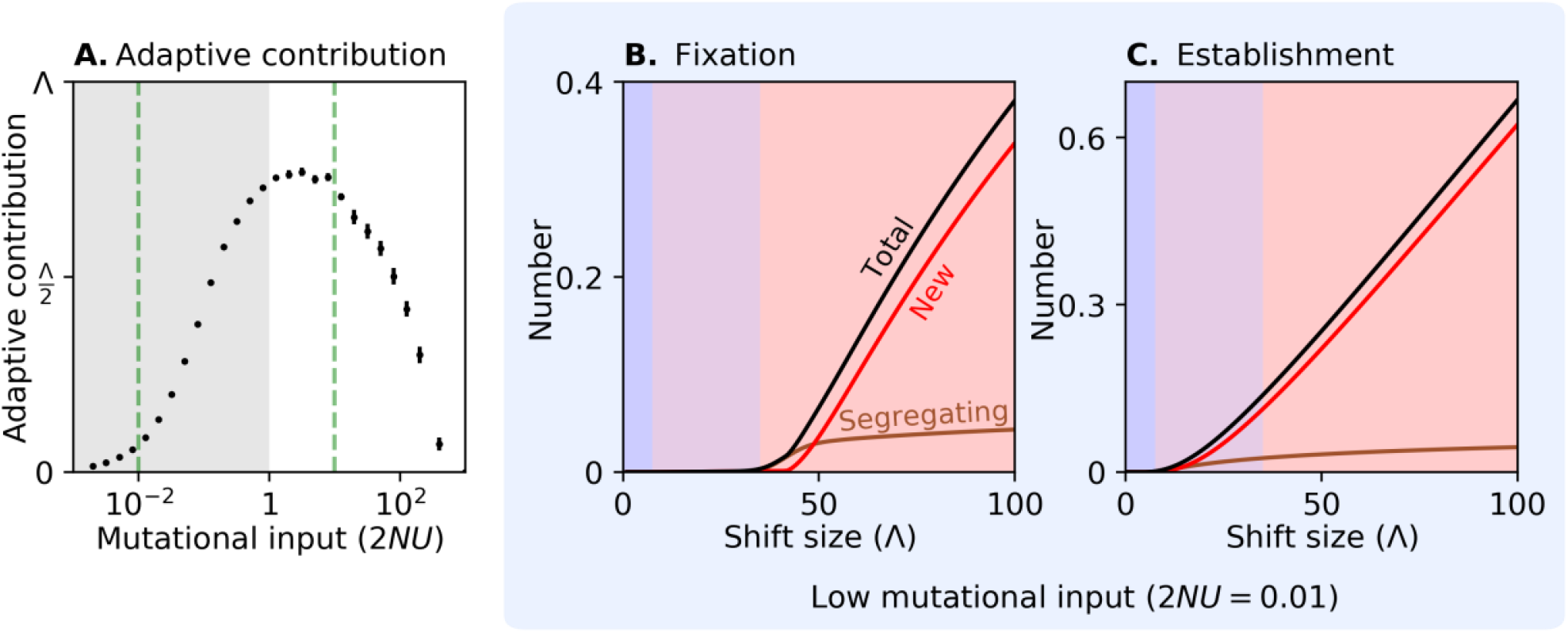
The long-term contribution of large effect alleles to adaptation. The mean contributions (±2*sD*) shown in (A) were calculated from 600 replicas of simulations with *N* = 5000, the distribution of effect sizes in Figure 1, Λ= 80 and *σ*^2^ = 40. The expected number of large effect fixations (in B) and large effect establishments (in C) were calculated assuming the same parameters as in (A), with 2*NU* = 0.01 (see supplement section 6.1). Simulation results closely match the analytic calculations (see Figure S7) and were omitted for clarity. Shaded regions in (B) and (C) correspond to regions defined as in Figure 3E.

When the mutational input increases further (i.e., when 2*NU* ≳ 1), however, the dependence on mutational input reverses. In this range, multiple large effect alleles establish themselves in the population and increase the rate of adaptation, which, in turn, shortens the duration of strong directional selection on each one of the alleles. This way, large effect alleles interfere with each other’s chances of fixing. The interference becomes stronger as the mutational input increases; when it is sufficiently large, interference prevents any large effect allele from fixing and their adaptive contribution zeroes out.

To gain a better sense of the underlying dynamics, we flesh out one example in each of these ranges (corresponding to the green dashes lines in Figure 4A). With typical human parameters, the low input example of 2*NU* = 0.01 corresponds to a target size of approximately 25 bp; this could occur, for instance, if a limited number of amino acid changes to a protein are highly beneficial. The high input example of 2*NU* = 10 corresponds to a target of approximately 25K bp, as might reflect a scenario in which loss of function (LoF) mutations in a few dozens of genes could drive substantial adaptation.

### Low mutational input (2*NU* < 1)

Figure 4B shows how the expected number of large effect alleles that fix depends on the shift size when the mutational input is low (supplement section 6.1). These results mirror the case with one large effect allele (Figure 3D and E). When shift sizes are too small, alleles cannot even establish themselves (in the blue region); when shifts exceed a certain size, alleles can establish themselves but cannot reach fixation (in the purple region); and, when shift sizes exceed some greater threshold, segregating alleles that start at sufficiently high frequencies can fix in the population but new alleles cannot (at the low end of the red region). Once shift sizes become even larger, however, most fixations arise from new mutations (most of the red region), because there are many fewer alleles that segregate before the shift than mutations that arise shortly enough after it (see Figure 4C, where the expected number of established alleles is separated into the contributions from segregating and new alleles).

### High mutational input (2*NU* / 1)

In the case with a high mutational input, many large effect alleles segregate at the time of the shift, markedly affecting the phenotypic dynamic. Figure 5A shows the mean distance of the population from the optimum as a function of time after the shift. Initially, the distance follows Lande’s approximation with a constant genetic variance (equal to the initial variance from both the Fisherian genetic background and large effect alleles). Shortly thereafter, many large effect alleles that are aligned with the shift rise in frequency, markedly increasing the genetic variance (Figure 5B) and accelerating phenotypic adaptation compared to Lande’s approximation. Not long after that, however, adaptation slows down markedly (note that the x-axis is on a log scale). This slowdown is caused by selection against individuals who carry several aligned, large effect alleles, whose phenotypes overshoot the new optimum (Figure 5B). This phenomenon is well understood (see the “non-Lande” case in Hayward and Sella 2022), so we do not explore it further here. Instead, we focus on the effects of these phenotypic dynamics on the trajectories of large effect alleles.

**Figure 5.**
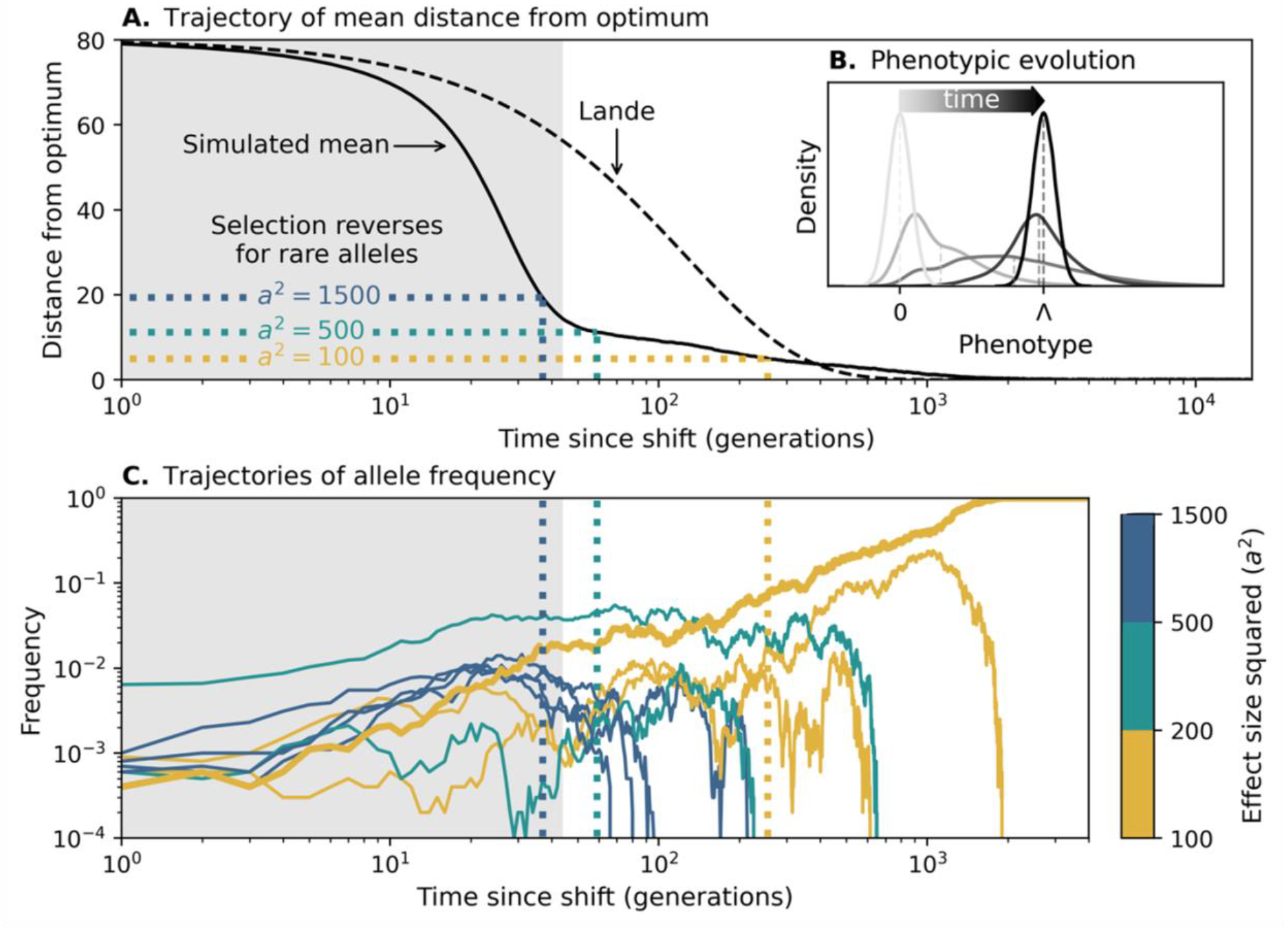
The adaptive response with many large effect alleles (for 2*NU* = 10). The mean distance from the optimum (shown in A) and phenotypic distributions (in B) are based on averaging 100 simulations with Λ= 80, *σ*^2^ = 40, *N* = 5000, and the distribution of effect sizes shown in Figure 1. The phenotypic distributions shown correspond to times: before the shift, 20, 42 (when adaptation slows down; see supplement section 6.2) and 300 generations, and after the population equilibrates around the new optimum. Large effect allele trajectories (shown in C) were randomly sampled from a single simulation with the same parameters to include 1-5 alleles that exceed frequency 1% and eventually go extinct in each bin of effect sizes (colored bars on the right), as well as one allele that fixes. The dotted lines that cross panels correspond to the distances and times where selection on rare alleles (*x* ≪ 1⁄2) with effect sizes *a*^2^ = 1500, 500 and 100 changes sign.

Figure 5C shows a representative sample of trajectories for large effect alleles that establish. These alleles are initially driven up in frequency by directional selection. However, selection on a rare allele with effect size *a* turns negative once the distance from the optimum drops below *a*/2, because the effect of stabilizing selection on it becomes stronger than the effect of directional selection (Eq. 1). This reversal of the sign of selection happens earlier for alleles with larger effect sizes. With many large effect alleles initially accelerating adaptation, those with the largest effects (in blue) get nowhere near frequency ½ before selection on them reverses, so they all eventually go extinct. Large effect alleles with intermediate effects (in teal) are positively selected for longer, allowing them to reach higher frequencies, but they still fall short of approaching frequency ½, and they also go extinct.

The large effect alleles with the smallest effects (gold) are positively selected for much longer, owing to the slowdown of phenotypic evolution. Those among them that make it to higher frequencies are positively selected for even longer, because the effects of stabilizing selection become weaker as they near frequency ½ (Eq. 1). When selection on these alleles eventually reverses, negative selection and genetic drift can have comparable effects on their frequencies for a while. Consequently, one of these alleles may cross the Rubicon at ½ and continue to fixation (thicker gold trajectory, with *a*^2^ = 110). Therefore, when interference allows large effect fixations, it favors alleles with smaller effect sizes (see supplement sections 6.2 and 6.3). Additionally, and in contrast to the case with low mutational input, interference favors the fixation of segregating alleles rather than new ones, because in this case, segregating alleles are abundant, and their head start gives them a competitive advantage (see Figure S6).

When the mutational input is sufficiently large, large effect alleles cannot fix at all (Figure 4A and supplement section 6.2). In this case, adaptation is spread over numerous large effect alleles, such that they all remain at low frequencies when adaptation slows down. Additionally, the slowdown occurs when the population mean is near the new optimum, such that even the smallest large effect alleles are selected against off the bat (see the “non-Lande” case in Hayward and Sella 2022).

### Putting the pieces together

Figure 6 shows how the probability that any large effect allele fixes depends on the main parameters of a trait (for the contribution to adaptation, see Figure S4). These dependencies mirrors what we found in previous sections. All else being equal, the probability of fixation and long-term contribution to adaptation increase with the mutation input when the input is low (2*NU* < 1), decrease with the mutational input when the input is high (2*NU* / 1), and are maximized when 2*NU*∼1 (Figures 6A, B, and S6). For a given mutation input, increasing the background genetic variance or decreasing the shift size reduces the time during which large effect alleles are positively selected and therefore reduces their probability of fixing and their long-term contribution to adaptation (Figures 6A, B and S6, respectively).

**Figure 6.**
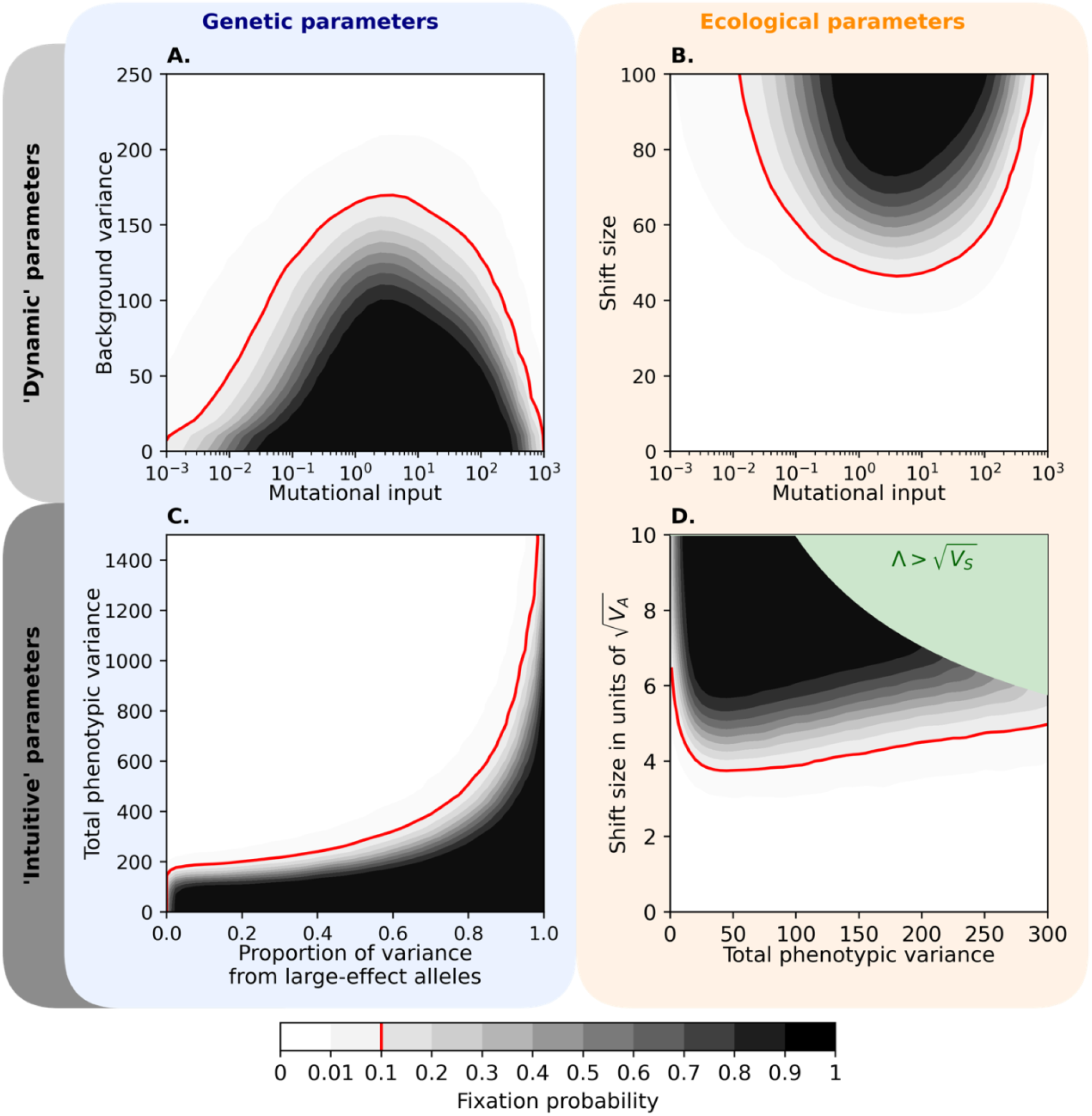
The probability of large effect fixations as a function of genetic and ecological parameters of traits. The contours were drawn by estimating fixation probabilities in simulations with an evenly spaced grid of parameter combinations and smoothing the results using a Gaussian noise filter (see supplement section 7.1 for more details). The model parameters were *N* = 5000, the distribution of effect sizes from Figure 1, Λ= 80 in (A) and (C), *σ*^2^ = 80 in (B), and a proportion of variance from large effect alleles *p* = 0.5 in (D). The green region in (D) is where 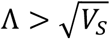(see supplement section 1). See Figures S13, S18, and S19 for alternative parameter values and the simulation results without smoothing.

### Relating our predictions with empirical findings

We can gain further insights by expressing trait parameters in terms of quantities that can be more readily estimated. In Figure 6C, we replace the key genetic parameters, *σ*^2^ and 2*NU* (in Figure 6A), with the total genetic variance in the trait at MSDB (before the shift), *V*_*A*_(0), and the proportion of the variance that arises from large effect alleles, *p*. The figure describes the adaptive response to a shift that is on the order of the width of the fitness function 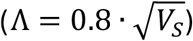 and entails an instantaneous mean fitness reduction of ∼27%. The red line denotes the threshold phenotypic variance that separates the two modes of adaptation: below it, the probability that large effect alleles reach fixation and contribute to long-term adaptation is appreciable; above it, long-term adaptation is dominated by the Fisherian genetic background. The threshold corresponds to the case in which the adaptive response is just slow enough to allow “lucky” large effect alleles to cross the Rubicon at frequency ½; it therefore increases with the size of the shift in optimum (see supplement section 7.2 and Figure S14). That the mode of adaptation changes—from a long-term response dominated by a few alleles to a highly polygenic one—as the total variance increases echoes results found under different settings (Lande 1983; Chevin and Hospital 2008; Höllinger et al. 2023; Sakamoto et al. 2024).

Data from human GWAS may allow us to infer where traits lie in Figure 6C, and thus whether we would expect large effect alleles to contribute to long-term adaptive changes in such traits. Several studies have estimated the proportion of phenotypic variance in traits that arises from common genetic variation (Yang et al. 2010; Yang et al. 2015; Speed et al. 2017; Karjalainen et al. 2024). If we assume, as is plausible, that common variation is dominated by variants with small and intermediate effects (per our definition) then these estimates suggest that the proportion of heritable variance explained by variants with large effects, *p*, for many traits is between 5-50% (supplement section 8.1). Simons et al. (2022) recently inferred the evolutionary parameters underlying genetic variation in 95 highly polygenic quantitative traits, including morphometric, molecular, and cardiovascular traits as well as blood phenotypes. Based on their results, we can estimate *n*_*e*_ · *V*_*A*_ for each trait, where *V*_*A*_ is the total genetic variance measured in our units and *n*_*e*_ is the ‘effective’ degree of pleiotropy. Here, *n*_*e*_ is defined as the number of independent traits, with the same association to fitness as the focal trait, that are required to account for the strength of selection acting on genetic variation affecting the focal trait (supplement section 1.2 in Simons et al. 2018). If *n*_*e*_ ≤ 3 then the total genetic variance in all 95 traits is greater than 300 (in our units); if *n*_*e*_ ≤ 45 than the same would be true for more than half the traits (supplement section 8.1, Figure S15). Taken together, these estimates suggest that many traits studied in human GWAS fall well above the red line in Figure 6C, i.e., that large effect alleles are not expected to contribute to the long-term adaptive response (although they are likely to have a transient contribution; see, e.g., Figure 3 and Figure S5). Improving these inferences (e.g., by using GWAS data for many traits jointly in order to estimate *n*_*e*_) should make it possible to predict the contribution of large effect fixations to the adaptive response to changes in the optima of many traits.

We can also relate our predictions with findings about large effect adaptative changes, by asking about the overlap between trait parameters under which we would predict such adaptive changes to occur and those for which commonly used methods to identify adaptive changes are well-powered. In supplement section 8.2, we consider a QTL study of a quantitative trait performed using a sample of 100 F2 hybrids between two closely related species. If the species split time is sufficiently recent and the contribution of small and intermediate alleles to variance in the trait is not negligibly small, our calculations suggest that such a QTL study should be well-powered to identify large effect adaptive changes throughout the trait range in which we predict they would occur (in Figure 6C). Similar analyses could be applied to footprints of selective sweeps in genetic polymorphism data, which reflect the specific trajectories of favored alleles (Barton 2000; Pennings and Hermisson 2006b; Pennings and Hermisson 2006a; Coop and Ralph 2012). As we illustrated in Figures 3 and 5 and in supplement section 5.3, our model can be used to predict these trajectories as a function of trait parameters and thus to delimit the trait range under which methods to identify sweeps would be well-powered.

These kinds of analysis should help us test our predictions and contextualize findings of large effect adaptive changes. Notably, finding examples of large effect adaptive changes in settings in which we would not predict them to occur would challenge our theory, whereas finding them only in regions where we would predict that they could occur would help validate it. Moreover, assessing the overlap between trait parameters in which current methods are well powered and in which we predict that large effect fixations could occur would help us evaluate how many large effect adaptive changes we are missing, and thus how common these changes are likely to be.

We can also ask about the likely contribution of large effect alleles as a function of shift size. To express shift sizes in more familiar terms, we measure them relative to the standard deviation of genetic variation in the trait, 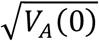(Figure 6D). When the genetic variance is tiny, shift sizes of several standard deviations are too small for large effect alleles to be beneficial immediately after the shift, let alone for them to reach fixation. Throughout the rest of the range, the shift sizes required for the fixation of large effect alleles is roughly proportional to 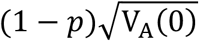(supplement section 7.2). For highly polygenic traits in humans, these shift sizes violate our condition that 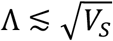. We imposed this condition not because of an inherent limitation of our analysis but rather because for larger shift sizes, we cannot assume that our fitness function approximates the true fitness effects. Moreover, enormous shifts in trait optima would plausibly result in a reduction in population size and even in extinction. These reservations notwithstanding, we can extrapolate our results to estimate what shift size in optimal human height, for example, would allow large effect alleles to fix. If we assume an *n*_*e*_ ≤ 3 for height then our bounds on genetic parameters suggest that this would require shift sizes exceeding 21 standard deviations, or roughly 1.4 meters (and this shift size scales with 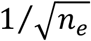; supplement section 8.1 and Eq. S62).

### Extensions to the genetics and ecology of traits

We studied the genetic basis of adaptation in a model of a panmictic population with a quantitative trait under stabilizing selection, which starts at MSDB before the trait optimum instantaneously shifts (Figure 1E). This model is rooted in classic, empirically-motivated assumptions. Moreover, it is general in the sense that adaptation from a few large effect changes and a highly polygenic response emerge within a single framework, allowing us to delimit the conditions under which we expect to observe a contribution of large effect alleles to adaptation. Nonetheless, our model makes restrictive assumptions about the genetics and ecology of a trait, which exclude factors that may affect the adaptive response.

Notably, given that genetic variation affecting one trait typically affects many others (Stearns 2010; Cotsapas et al. 2011; Sivakumaran et al. 2011; Lee et al. 2012; Andreassen et al. 2013; Bulik-Sullivan et al. 2015; Pickrell et al. 2016; Frei et al. 2019; Zhang 2023), we would like to know how such pleiotropic effects influence the adaptive response. One way to account for these is to model adaptation in a multidimensional trait space, akin to Fisher’s geometric model (Fisher 1930), where the optimal phenotype exhibits a shift in a “focal” trait while remaining fixed for the other traits. In this model, the effects of directional selection on alleles reflect selection on the focal trait, as they do in the current model, but the effects of stabilizing selection on alleles would be stronger, because they would absorb selection on all traits. The adaptive response from small and intermediate effect alleles would follow Lande’s multidimensional approximation (Lande 1976). For large effect alleles to fix, the net selection acting on them—including the negative selection arising from pleiotropic stabilizing selection—would have to remain positive and strong for long enough for them to cross the Rubicon at frequency ½.

In other words, for large effect alleles to fix, they would have to be fairly trait-specific, with appreciable (aligned) effects on the focal trait and minimal effects on other traits. As a result, pleiotropic effects reduce the reservoir of large effect alleles that can reach fixation (Fisher 1930; Kimura 1983; Orr 1998; Otto 2004; Chevin et al. 2010), an effect that can be loosely viewed as lowering the “effective” mutational input of large effect alleles (in, e.g., Figures 6A and B) and their “effective” contribution to variance (in Figure 6C). We might expect suitable large effect alleles to arise in genes and/or regulatory elements that are relatively specific to a trait (Spence et al. 2024). In such cases, the limited reservoir would cause fixations to reoccur in the same genes or elements under similar selective pressures (see, e.g., Stern 2013, for similar arguments). Consistent with these predictions, this kind of parallel evolution is observed for many instances of large effect adaptive changes (including all but one of the examples [EDA] in the Introduction).

Another interesting extension to the genetics of a trait in our model arises if we allow a high population-scaled mutation rate to large effect beneficial mutations at a single (tightly linked) locus. This might occur, for example, when LoF mutations to a gene are adaptive and/or when the population size is sufficiently large. In this case, multiple beneficial large effect alleles at a single locus might ascend in frequency after the shift in optimum. If we assume, for example, that these mutations have the same effects on the focal trait, then we can understand their long-term contribution to adaptation by considering all of them as a single large effect allele in our model. Namely, if their aggregate frequency reaches frequency ½ before the population mean nears the new optimum, these mutations would eventually replace the ancestral allele. After they do, random genetic drift will eventually drive one of them to fixation, but until this occurs, multiple large effect alleles would segregate in the population. This kind of partial sweep was described by Pennings and Hermisson (2006) and Coop and Ralph (2012) in the case of directional selection at a single locus; Ralph and Coop (2010) also considered how such sweeps play out within a spatially extended range. This kind of partial sweep in space might explain cases in which several large effect adaptive changes at the same locus are found segregating within a species range, including multiple LoF mutations in *FRIGIDA* found segregating in *Arabidopsis thaliana* in response to selection on flowering time (Johanson et al. 2000; Le Corre et al. 2002; Shindo et al. 2005); independent mutations in a *cis*-regulatory element that reduce pigmentation in *Drosophia santomea* (Jeong et al. 2008); recurrent deletions of a *PitX* enhancer that causes pelvic reduction in sticklebacks (Chan et al. 2010); and multiple examples in humans including lactase persistence (reviewed in Novembre and Di Rienzo 2009).

We would also like to extend the model to consider additional aspects of a trait’s ecology, notably variation in selection pressures on a trait over space and time. Temporal variation in selection pressures, ranging from seasonal to glacial cycles (e.g, Lister 2004; Willis et al. 2004; Bergland et al. 2014; Enbody et al. 2023), affects the genetic architecture of a trait. In particular, minor allele frequencies and heritable variance across the range of allele effect sizes may increase, and in certain conditions, balancing selection may maintain stable, large effect polymorphisms (Haldane and Jayakar 1963; Gillespie 1973; Bürger and Gimelfarb 2002; Wittmann et al. 2017). Changes to genetic architecture could markedly affect the response to selection, and specifically the probability of large effect fixations.

Many examples of large effect adaptations involve selection pressures that vary over a species range, whose evolution was plausibly accompanied by migration between different local environments. Alleles that are beneficial in one environment and detrimental in another may spread and be maintained at migration-selection balance—a form of balancing selection—if their selection effects are large relative to rates of migration and the effects of genetics drift (Levene 1953; Bulmer 1972; Yeaman and Otto 2011, reviewed in Lenormand 2002; Yeaman 2022). This theory could help explain why LoF mutations in the *FRIGIDA* gene in *Arabidopsis thaliana* was favored as an adaptation for flowering time (Le Corre et al. 2002; Stinchcombe et al. 2004; Toomajian et al. 2006), and why an extremely old LoF allele in EDA present at low frequency in marine environments, has repeatedly contributed to armor plate reduction in freshwater habitats (Colosimo et al. 2005; Jones et al. 2012; reviewed in Reid et al. 2021). Theoretical studies have also described the conditions for the spread and maintenance of locally adapted ‘supergenes’ (Kirkpatrick and Barton 2006; Yeaman and Whitlock 2011; Yeaman 2013), i.e., sets of multiple locally adapted alleles held together in an inversion or an otherwise tightly linked locus, and several examples of such supergenes have been identified in recent years (reviewed in Schwander et al. 2014; Gutiérrez-Valencia et al. 2021). Theoretical studies of the evolution of polygenic, quantitative traits with optima that vary among habitats suggest that in principle large effect alleles or ‘supergenes’ could be favored and maintained in this setting, but only so long as evolutionary parameters fall within specific ranges (reviewed in Yeaman 2022; see also Sakamoto et al. 2024). These studies explored simple scenarios, e.g., of a two-island model with constant selection and migration, when spatiotemporal variation in selection and migration can take many forms. The question about when and how often the interplay between spatiotemporally varying selection, migration, and drift allow large effect alleles to spread and be maintained therefore remains largely open.

### Outlook

In summary, our model makes clear that the contributions of large effect and polygenic changes to adaptation depend on genetic and ecological properties of traits. By making these relationships explicit, we can start to predict what we expect to find based on trait parameters, relate these predictions with disparate empirical findings, and begin to answer enduring questions about the genetic basis of adaptation.

## Acknowledgements

We thank Peter Andolfatto, Andrés Bendesky, Jiarun Chen, Graham Coop, Hannah Munby, Magnus Nordborg, Jonathan Pritchard, and Molly Przeworski for many helpful discussions, Yuval Simons for estimates of variance in human traits, and Molly Przeworski for comments on the manuscript. This work was supported by NSF GRFP grant DGE1644869 to WRM and NIH R01 grant GM115889 to GS.

