## Supplement Note for "When should adaptation arise from a polygenic response versus few large effect changes?"

### Supplementary Information

William R Milligan[1,*], Laura K Hayward[2] and Guy Sella[1,3,*]

[1] Department of Biological Sciences, Columbia University, New York, NY, USA

[2] Institute of Science and Technology Austria, Klosterneuburg, Austria

[3] Program for Mathematical Genomics, Columbia University, New York, NY, USA

### List of Tables

Table S1. Summary of notation. ... 3
Table S2. Assumptions on model parameters. ... 6
Table S3. Estimates of heritability explained by common genetic variation. ... 38

### List of Figures

Figure S1. The number of large effect fixations in individual- and allele-based simulations. 8
Figure S2. Contribution of small and intermediate effect alleles to phenotypic variance and skew. ... 9
Figure S3. Polygenic adaptation causes small changes to fixation probabilities at MSDB. . 11
Figure S4. Comparison of approximations of the fixation probability for (A) segregating and (B) new alleles ... 16
Figure S5. Allelic and phenotypic trajectories. ... 21
Figure S6. Long-term adaptive response of large effect alleles vs. their mutational input. . 23
Figure S7. Number of alleles that fix with low mutational input. ... 26
Figure S8. The time (A) and distance (B) when the quasistatic adaptation phase begins. .. 29
Figure S9. Effect size distributions of fixed alleles for a tiny mutational input. ... 30
Figure S10. Effect size distributions of fixed alleles. ... 31
Figure S11. The genetic basis of adaptation with midrange effect alleles. ... 32
Figure S12. Probability of fixation for single aligned, segregating allele vs effect size. ... 34
Figure S13. Phase spaces without smoothing corresponding to Figure 6. ... 35
Figure S14. Simple approximations for parameter ranges allowing for fixation. ... 37
Figure S15. Estimates of genetic variance for the 95 traits from Simons et al. (2022). ... 40
Figure S16. The power to identify a large effect fixation in a QTL study. ... 43
Figure S17. The adaptive contribution of large effect fixations as a function of the genetic and ecological parameters of traits. ... 44
Figure S18. Alternate choices for fixed parameters in Figure 6. ... 45
Figure S19. Alternate distribution of selection effects in Figure 6. ... 46

**Table S1. Summary of notation.**

| Parameter | Meaning |
|---|---|
| $N$ | Population size |
| $V_S$ | The width of the Gaussian fitness function |
| $\delta = \sqrt{V_S/2N}$ | The units we use to measure the trait and allelic effect sizes |
| $t$ | Time since shift in generations |
| $U$ ($U^*$) | Mutational input of large effect alleles (of all alleles) per gamete per generation |
| $\Lambda$ | Magnitude of shift in trait optimum |
| $\mu$ | Mean phenotype in the population |
| $D(t)$ | Difference between mean phenotype and trait optimum |
| $D_L(t)$ | Lande's approximation for $D(t)$ |
| $V_A(t)$ | Additive genetic variance, we denote the variance at MSBD as $V_A(0)$ |
| $\sigma^2$ | Genetic variance from small & intermediate effect alleles |
| $p$ | Proportion of variance from large effect alleles |
| $\mu_3(t)$ | Third central moment |
| $W(z)$ | Fitness of individuals with trait value $z$ |
| $\overline{W_\iota}(a)$ | Mean fitness of individuals with $i$ copies of an allele with effect size $a$ |
| $\overline{W}$ | Mean fitness of the population |
| $a$ | Allele effect size |
| $g(a)$ $\left(g^*(a)\right)$ | Distribution of effect sizes for large effect alleles (for all alleles) |
| $g_f(a)$ | Distribution of fixed effect sizes for large effect alleles |
| $\hat{g}(s)$ | Estimated distribution of selection coefficients |
| $a_{min}$ | Minimum effect size included in $g(a)$ |
| $s_0$ | Initial selection coefficient at the time of the shift or when it first arises, whichever is later |
| $s_e$ ($S_e$) | (Scaled) selection coefficient at MSDB |
| $s_d$ | Selection coefficient arising from directional selection |
| $x(t)$ | Allele frequency |
| $x_0$ | Initial allele frequency at the time of the shift or when it first arises, whichever is later |

| | |
|---|---|
| $x_{msdb}$ | Scalar that describes the distribution of frequencies for large effect alleles at MSDB |
| $x_{est}$ $(t_{est})$ | Scalar that describes the frequencies that allow for segregating alleles to establish (times for new alleles). |
| $x_c$ $(t_c)$ | Frequency (time) that allows large effect alleles to fix |
| $\tau_{msdb}(x\|a)$ | Sojourn time of alleles at frequency $x$ given $a$ at MSDB |
| $\tau(x)$ $\left(\tau(x\|x_\infty = x^*)\right)$ | Sojourn time of alleles at frequency $x$ after the shift (conditional on absorption at boundary $x^*$). |
| $p_{msdb}(x\|a)$ | Density of alleles at frequency $x$ given $a$ at MSDB |
| $P_s(fix)$ $\left(P_n(fix)\right)$ | Probability that a single segregating (or new) allele reaches fixation |
| $P_s(est)$ $\left(P_n(est)\right)$ | Probability that a single segregating (or new) allele establishes |
| $n_{f\|s}$ $(n_{f\|n})$ | Number of segregating (or new) alleles that fix |
| $n_{e\|s}$ $(n_{e\|n})$ | Number of segregating (or new) alleles that establish |
| $t_{1/2}$ | Time at which a given allele reaches frequency ½ |
| $t_{qs}$ | Time at which the quasi-static phase begins |
| $D_{qs}$ | Distance at $t = t_{qs}$ |
| $h_q^2$ | Heritability arising from variants with minor allele frequency greater than $q$ |
| $n$ | Pleiotropic factor |
| $\eta$ | Sample size of QTL study |
| $h_L^2$ $(V_L)$ | Heritability (variance) contributed by a large effect fixation |
| $\mathcal{P}$ | Power to detect a large effect fixation in a QTL study |
| $V_T$ | Total variance in a hybrid population |
| $V_E$ | Environmental contribution to phenotypic variance |
| $V_{pol}$ $(V_{div})$ | Phenotypic variance in a F2 hybrid population contributed by alleles segregating (or differentially fixed) in the two parent populations |

### 1. Assumptions on parameter ranges

The scope of our investigation is defined in terms of several conditions on the parameters of the model. These conditions are summarized in Table S2.

The first set of conditions ensure that the phenotypic distribution at mutation-selection-drift balance (MSDB) is tightly centered around the optimal trait value and that genetic variation at MSDB is not subject to extremely strong selection. First, we assume that the phenotypic variance at MSDB is much smaller than the width of the Gaussian fitness function, *i.e.*, that $V_S \gg V_A(0)$. This assumption ensures that, when the population mean phenotype is near the optimum, individual phenotypes are near the trait optimum and that our results are insensitive to the shape of the fitness function, which we generally expect to be quadratic near an optimum but do not have a general expectation for far from an optimum. The assumption on $V_A(0)$ translates into a condition on the total rate of mutations affecting the trait. Specifically, the expected contribution to phenotypic variance per unit mutational input is bound by $5\delta^2$ (see, *e.g.*, Fig. 2 in Simons et al. 2018) so we require that the total mutation rate per gamete per generation satisfies $U^* < 0.02$. Given mutation rates per site per generation are typically very small (approximately $1.25 \cdot 10^{-8}$ in humans, see *e.g.*, Kong et al. 2012), this still allows for large mutational target sizes (up to 1.6 Mb in humans). Second, to ensure that the phenotypic distribution at MSDB is centered at the optimal trait value we require that the mutational distribution of effect sizes be symmetric, *i.e.*, that $g^*(a) = g^*(-a)$. Third, we make the standard population genetic assumption that the selection coefficients of alleles satisfy $s \ll 1$. At MSDB, the selection coefficient of an allele with effect size $a$ is $s_e \approx a^2/V_S$ (Wright 1931; Wright 1935; Turelli 1984), so we require that $a^2 \ll V_S$. Fourth, we assume that a substantial proportion of new mutations are not effectively neutral, $E(a^2) \gtrsim 1$, which ensures that a selective response to the shift in the optimum can be observed. Under these assumptions, the mean phenotype exhibits tiny fluctuations around the optimal phenotype with variance $\delta^2 = V_S/2N$ (Simons et al. 2018). We define our units such that $\delta^2 = 1$, which in turn defines $V_S = 2N$ and $S_e = 2Ns_e = a^2$ at steady state.

The second set of conditions ensure that selection on individuals and variants after the shift in optimum is not extremely strong. To this end, we assume that the shift size does not exceed the width of the fitness function, *i.e.*, that $\Lambda \leq \sqrt{V_S}$. Like our condition on phenotypic variance at MSDB, this condition ensures that our results are insensitive to the shape of the fitness function. It also ensures that the reduction in mean population fitness (the genetic load) is not too large, which makes our assumption that the population size remains constant sensible. This assumption still allows for shifts that are several times greater than the phenotypic standard deviation at MSDB (because $V_A(0) \ll V_S$).

We also assume that the directional selection coefficients of alleles after the shift satisfy $s_d \ll 1$. The directional selection coefficient is well approximated by $s_d \approx aD/V_S$ (Wright 1931; Barton and Turelli 1987; Hayward and Sella 2022). Our previous assumptions that $a^2 \ll V_S$ and $D \leq \Lambda \leq \sqrt{V_S}$ already imply that $s_d < 1/\sqrt{10}$. With large shift sizes ($\Lambda \approx \sqrt{V_S}$) and effect sizes, we could still get $s_d > 0.1$ immediately after the shift, but this deviations from our assumption would be short lived unless the phenotypic variance is minute. Below, we show that our results are insensitive to this kind of strong selection (Fig. S1), so we refrain from adding additional conditions on parameters to exclude these cases.

**Table S2. Assumptions on model parameters.**

| Assumption | Interpretation |
| --- | --- |
| $U^* < 0.02$ ($V_A \ll V_S$) | Phenotypic variance is much smaller than the width of the fitness function |
| $g^*(a) = g^*(-a)$ | Symmetric mutation |
| $a^2 \ll \sqrt{V_S}$ | Allele effect sizes are much smaller than the width of the fitness function |
| $E(a^2) \gtrsim 1$ | A substantial proportion of alleles are not effectively neutral |
| $\Lambda \leq \sqrt{V_S}$ | Shift sizes are not massive relative to the width of the fitness function |

### 2. The allelic and phenotypic equations

The allelic and phenotypic equations that we use were derived in Appendix 3 of Hayward and Sella (2022). Here we recount the main steps in these derivations, as well as a specific simplification that we employ.

The derivation starts from the standard equation for the expected change in allele frequency in a single generation. For an allele with effect size $a$ at frequency $x$ this equation takes the form

$$E(\Delta x) = E(x') - E(x) = \frac{x^2\bar{W}_2(a) + x(1-x)\bar{W}_1(a)}{\bar{W}}, \quad \text{(S1)}$$

where $\bar{W}_i(a)$ is the mean fitness of individuals carrying $i$ copies of the focal allele and $\bar{W}$ is the mean fitness of the population. The mean fitness of individuals carrying $i$ copies of the focal allele is then approximated by

$$\bar{W}_i(a) \approx \int_R W(R + ia) f(R) dR \approx W\big(\mu + a(i - 2x)\big), \quad \text{(S2)}$$

where $f(R)$ denotes the distribution of background genetic contributions to the phenotype, $R$, and $\mu$ is the mean phenotype in the population. This approximation assumes that the background contribution is independent on the focal genotype, which is sensible given our assumption of free recombination. It also neglects the effects of variation in the background contributions, which is sensible given our assumption that $V_A \ll V_S$. Under this approximation, a second-order Taylor expansion around $a/\sqrt{V_S} = 0$ yields the allelic equation in Hayward and Sella (2022):

$$E(\Delta x) \approx x(1-x) \cdot \frac{a}{V_S}\left(D(t) - a\left(1 - \frac{D^2(t)}{V_S}\right)\left(\frac{1}{2} - x\right)\right). \tag{S3}$$

We use a simplified version of this equation (Eq. 1) that neglects $D^2(t)/V_S$, which provides a decent approximation except for the period immediately after a large shift in optimum (*i.e.*, $\Lambda \approx \sqrt{V_S}$).

To derive the phenotypic equation, Hayward and Sella (2022) sum over all the allelic equations. This way they find that

$$E(\Delta \mathrm{D}) \approx -V_A(t) \cdot \frac{D(t)}{V_S} - \frac{\mu_3(t)}{2V_S}\left(1 - \frac{D^2(t)}{V_S}\right), \tag{S4}$$

where $\mu_3(t)$ is the third central moment. Here too, we neglect the term $D^2(t)/V_S$.

### 3. Simulations

#### 3.1. Justifying the use of simplified simulations

Throughout the manuscript, we use simulations that realize a simplified version of our model. Rather than realizing the full model that is described in terms of a population of individuals (Fig. 1D), we model the dynamics of large effect alleles in terms of their allelic equations (Eq. 1), whereas the effects of small and intermediate effect alleles are approximated in terms of Lande's approximation (Eq. 3) assuming a constant contribution to variance, $\sigma^2$. Here we justify the use of these simplified simulations.

First, we consider the approximation of the dynamics in terms of alleles rather than individuals. We test the accuracy of this approximation in what we consider our two worst case scenarios: 1) when the trait's genetic variance is very small ($V_A(0) \approx 1$) and the shift size is large such that large effect alleles may experience prolonged strong selection with selection coefficients above 0.1, 2) when the trait's genetic variance is very large ($V_A(0) \approx V_S/2$) such that the approximation used in Eq. S2 is not accurate. In Fig. S1, we show that simulations that track individuals and those that track alleles generate similar results in terms of the probability of a large effect fixation in both of the scenarios (see Hayward and Sella (2022) for description of simulations). For the rest of the parameter space, we rely on the results from Hayward and Sella (2022), which established the accuracy of this approximation in the highly polygenic case (when $V_A(0) \gg 1$). By simulating the dynamics in terms of alleles rather than individuals, we greatly improve the computational efficiency of our simulations without compromising accuracy.

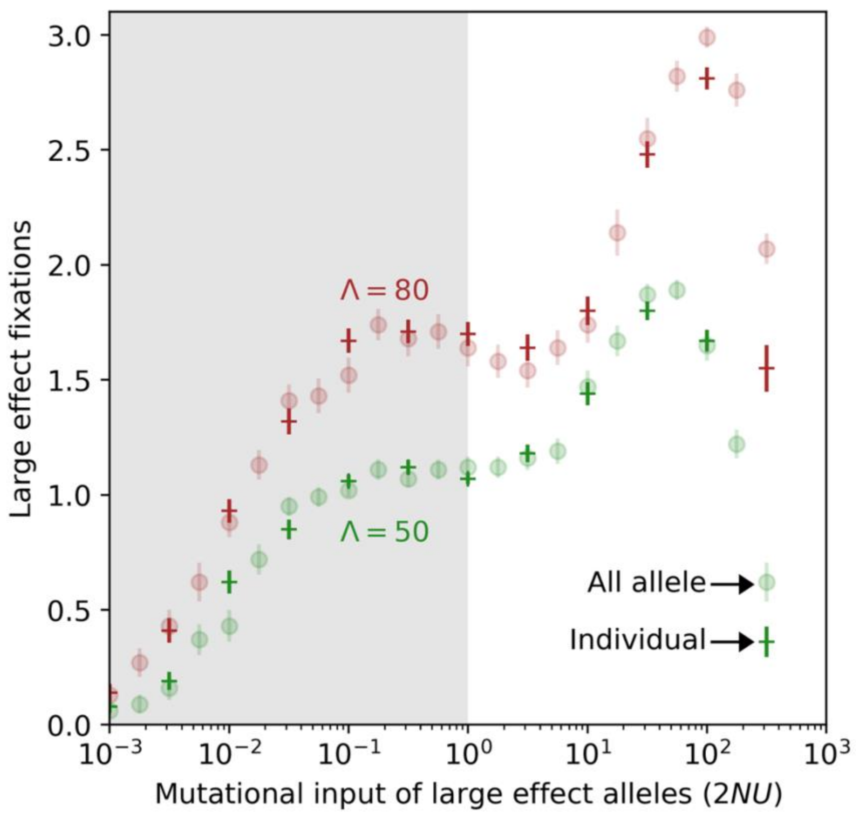

**Figure S1. The number of large effect fixations in individual- and allele-based simulations.** We assumed a mixture of two effect size distributions: an exponential distribution with a mean of $a^2 = 1$ for small and intermediate effect alleles and the exponential distribution described in Fig. 1 for large effect alleles. We chose the total mutational input (for all alleles) and the corresponding weights for the effect size distributions such that the expected contribution of small effect and intermediate effect alleles to variance at MSDB equals $\sigma^2 = 4$ and the mutational input of large effect alleles equals the prescribed $2NU$ (on the x-axis). Each point represents the mean $\pm$2SEs calculated based on 100 simulations.

Second, we show that the effect of small and intermediate effect alleles is well approximated in terms of an infinitesimal genetic background. That is, the variance contributed by small and intermediate effect alleles is approximately constant and such alleles introduce negligible skew in the phenotypic distribution. Again, Hayward and Sella (2022) demonstrated the accuracy of this approximation in the highly polygenic case. In Fig. S2, we show the accuracy of this approximation for different combinations of shift size, background variance (here, the variance contributed by small and intermediate effect alleles), and mutational input of large effect alleles. For all combinations, the contribution of small and intermediate effect alleles to phenotypic variance remained within 30% of its value at MSDB with the most substantial deviations arising late in the adaptive process when the background variance is small. In contrast, the contribution of large effect alleles to phenotypic variance can change by orders of magnitude during the adaptive process. Additionally, for all parameter combinations, the contribution of small and intermediate effect alleles to phenotypic skew remained negligibly small and incapable of slowing adaptation while the population mean phenotype is far from the optimum. Therefore, we approximate the contribution of small and intermediate effect alleles to adaptation as the contribution from an infinitesimal genetic background and only explicitly simulate the trajectories of large effect alleles.

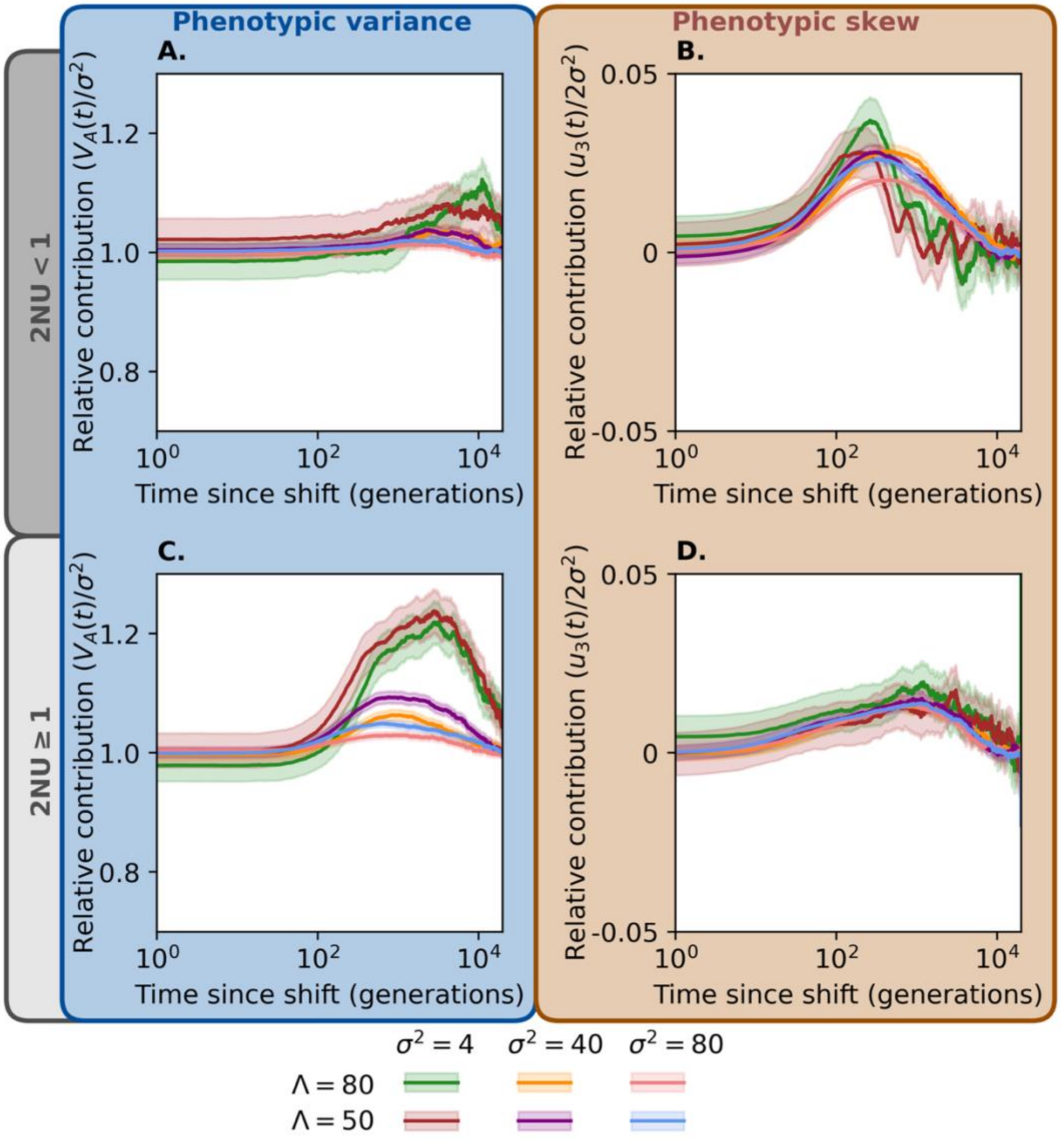

**Figure S2. Contribution of small and intermediate effect alleles to phenotypic variance and skew.** We show the contributions to phenotypic variance (A and C) and the third central moment (B and D) from alleles with $a^2 < 100$ as a function of time after the shift. The phenotypic variance is scaled relative to its value at MSDB. The third central moment is scaled relative to $2\sigma^2$ and thus represents how close the population mean phenotype must be to the new optimum for the population to enter the quasi-static phase at any given time; values much smaller than one indicate phenotypic skew has negligible effects on the adaptive dynamics. We simulated 100 replicates with the effect size distributions described in Fig. 1 for each combination of shift size, background variance, and mutational input ($2NU$). We used 6 logarithmically spaced values of $2NU$ between 0.001 and 100. Each curve shows the average and $\pm 2$ standard errors across replicates and grouped values of $2NU$.

#### 3.2. Description of simulations

We use two types of simulations, *simplified all-allele* simulations that implement our general model and *single large effect allele* simulations that track the trajectory of a single large effect allele. Both types of simulations make use of the approximations described in 3.1. Here, the frequency of large effect alleles in generation $t+1, x_{t+1}$, after drift and selection in generation $t$ is simulated using the following binomial distribution

$$x_{t+1} \sim Bin\big(2N, x_t + E(\Delta x_t)\big), \tag{S5}$$

where $E(\Delta x_t)$ is approximated from Eq. 1. The genetic background of small and intermediate effect alleles only affects the change in the distance between the mean phenotype and the optimal trait value each generation, which becomes

$$\Delta D(t) = -\underbrace{\textstyle\sum_{i \in L} 2a_i \Delta x_i(t)}_{\substack{\text{change due to} \\ \text{large effect alleles}}} - \underbrace{\frac{\sigma^2 D(t)}{V_S}}_{\substack{\text{change due to} \\ \text{genetic} \\ \text{background}}} . \tag{S6}$$

The first kind of simulation, *simplified all-allele*, allows large effect alleles to arise each generation. We initialize these simulations by the expected number of segregating sites and distribution of allele frequencies and effect sizes given the model parameters (see Hayward and Sella (2022) for the derivation of these quantities). Initializing in this way allows us to not include a burn-in period, which further increases computational efficiency. After initialization, the number of new alleles generated each generation is Poisson distributed with mean $2NU$ and their effect sizes are drawn from the given distribution, $g(a)$. These simulations are run until the population phenotypic distribution returns to MSDB around the new optimum and for a minimum of $4N$ generations.

The second kind—*single large effect allele simulations*—is similar to those in Chevin and Hospital (2008), which we use for illustrative purposes. These simulations assume there is a single large effect allele with a given effect size that is either segregating at the time of the shift or arises after the shift. When simulating a segregating allele, the initial frequency at the time of the shift is drawn from the stationary distribution at steady state. When simulating a new allele, the time that it arises is drawn from a uniform distribution between $t = 0$ and $t = \ln(\Lambda) V_S/\sigma^2$, the expected time for the population mean phenotype to be $\delta$ far from the new optimum with no contribution from large effect alleles. These simulations are run until the large effect allele reaches fixation or goes extinct, after which the simulations become deterministic.

Code to run simulations and generate figures can be found at https://github.com/wm2377/LargeEffectAdaptation.

**4. The conditions for the highly polygenic case**

We follow Hayward and Sella (2022) in describing the adaptive response in the highly polygenic case. In addition to the conditions on parameters that we detailed in section 1, Hayward and Sella (2022) make two additional assumptions to ensure that the adaptive response is highly polygenic. First, they assume the trait is highly polygenic, *i.e.*, that $2NU^* \gg 1$. Second, they assume that only a small average frequency change per segregating site is required for the population mean to reach the new optimum. The latter condition translates into the requirement that $\Lambda/\sqrt{V_A(0)} \lesssim \frac{1}{2}\sqrt{2NU}$. Under these conditions, adaptation is generally achieved by small (relative) changes to the fixation probabilities of alleles at MSDB, with a slight increase for alleles whose effects are aligned with the shift and decrease for alleles with opposing effects (see Fig. S3).

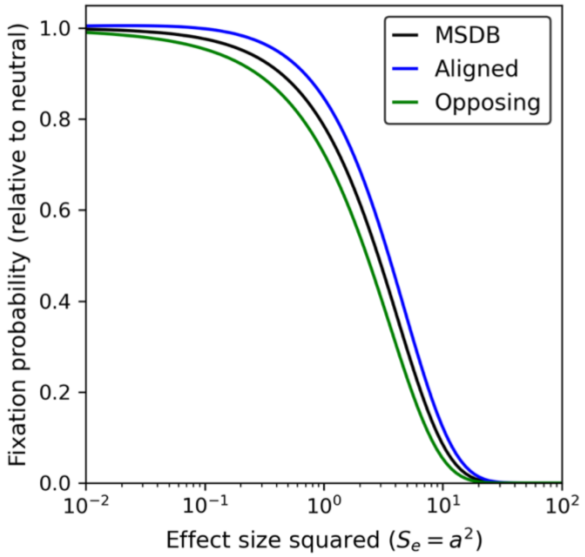

**Figure S3. Polygenic adaptation perturb fixation probabilities at MSDB.** As an illustration, we compare the fixation probability as a function of allele squared effect sizes with shift size

$\Lambda = 40$ and genetic variance $V_A = 400$ (calculated using the linear approximation in Hayward and Sella (2022)) with the fixation probabilities at MSDB. Fixation probabilities are scaled relative to those for neutral mutations, which renders them insensitive to the population size.

### 5. One large effect allele and a 'Fisherian' background

When we consider the case of one large effect allele and a 'Fisherian' background, we explain why the probability that the allele fixes can be approximated as the probability that it establishes itself in the population *multiplied* by the probability that it reaches frequency ½ conditional on establishing. In the first part of this section we derive such approximations for both segregating and new alleles. When an allele fixes or is lost it may do so following different kinds of trajectories. For example, it may be propelled all the way to fixation by positive directional selection resulting in a full selective sweep, or alternatively, directional selection on it may reverse after it crosses the Rubicon at frequency ½ resulting is a kind of partial sweep. In the second part of this section, we characterize these trajectories and drive the conditions in which they arise.

#### 5.1. The fixation probability of a segregating allele

**The probability of establishment.** The establishment or loss of the allele occur quickly after the shift in optimum, so we approximate the probability of establishment based on the selection acting on the allele immediately after the shift. In particular, we approximate this probability for an allele with effect size $a$ and initial frequency $x_0$ using Kimura's formula for the probability of fixation with constant additive selection (Crow and Kimura 1970):

$$P_s(\mathrm{est}|x_0, a) \approx \frac{1 - \mathrm{Exp}(-4Ns_0x_0)}{1 - \mathrm{Exp}(-4Ns_0)}, \tag{S7}$$

with the selection coefficient

$$s_0 = \frac{a}{V_S}\big(\Lambda - a(1/2 - x_0)\big), \tag{S8}$$

which includes both the directional and stabilizing selection terms immediately after the shift (Eq. 1). Given that the allele is initially rare, *i.e.*, $x_0 \ll 1$, its initial scaled selection coefficient is well approximated by

$$2Ns_0 \approx a(\Lambda - a/2). \tag{S9}$$

For the allele to establish with appreciable probability, the selection favoring it must be strong relative to genetic drift, *i.e.*, $2Ns_0 \gg 1$. Intuitively, this condition amounts to the effect of directional selection after the shift being substantially greater than that of stabilizing selection. In this case, the approximation for the probability of establishment simplifies to

$$P_s(\text{est}|x_0, a) \approx 1 - \text{Exp}(-2a(\Lambda - a/2)x_0) = 1 - \text{Exp}(-x_0/x_{est}), \tag{S10}$$

with

$$x_{est} \equiv 1/\big(2a(\Lambda - a/2)\big). \tag{S11}$$

We approximate the probability that an allele segregating before the shift fixes—without conditioning on the initial frequency $x_0$—by integrating over the distribution of initial frequencies before the shift, at MSDB. The probability density of the frequency of a (minor) large effect allele at MSDB is well approximated by

$$\rho_{msdb}(x_0|a) \approx C \cdot \text{Exp}(-a^2 x_0)/x_0 = C \cdot \text{Exp}(-x_0/x_{msdb})/x_0, \tag{S12}$$

where $C$ is a normalization constant such that $\int_0^{1/2} \rho_{msdb}(x_0|a)dx_0 = 1$ and

$$x_{msdb} \equiv 1/a^2 \tag{S13}$$

is the scale of allele frequency at MSDB (Appendix 3 in Hayward and Sella (2022)). This way we find that the probability that an allele with effect size $a$ establishes

$$\begin{aligned} P_s(\text{est}|a) &\approx \int_0^{1/2} P_s(\text{est}|x_0, a) \cdot \rho_{msdb}(x_0|a)dx_0 \\ &\approx C\left(\text{Ln}\left(1 + \frac{x_{msdb}}{x_{est}}\right) + h\left(\frac{1}{2x_{msdb}} + \frac{1}{2x_{est}}\right) - h\left(\frac{1}{2x_{msdb}}\right)\right) \end{aligned} \tag{S14}$$

where we assumed that $x_{est} > 0$, or equivalently, that $\Lambda < a/2$ (otherwise the allele would never establish) and where $h(y) \equiv -Ei(-y)$ and $Ei(y) \equiv \int_{-\infty}^{y} \text{Exp}\,(t)/t\,dt$ is the exponential integral. For large effect alleles ($a^2 \gg 1$), we can neglect the second and third terms in the integral's solution because $C \cdot h(y) \ll 0.1$ when $y \geq 5$. Thus, the probability of establishment for a segregating allele with effect size $a$ is well approximated by

$$P_s(\text{est}|a) \approx C \cdot \text{Ln}(1 + x_{msdb}/x_{est}) = C \cdot \text{Ln}(2\Lambda/a). \tag{S15}$$

**The probability of reaching frequency ½.** Next, we consider the probability that a segregating large-effect allele makes it to frequency ½ conditional on establishing. If we neglect the stochasticity in the allele's rate of ascent at low frequencies (*e.g.*, during establishment), we expect there to be some critical initial frequency, $x_c$, such that an allele that starts at this frequency and establishes will reach frequency ½ when the distance to the optimum hits 0. Alleles that start above this critical frequency will eventually fix, whereas those that start below it will be lost. We can therefore approximate the probability that a segregating allele with effect size $a$ fixes by

$$P_s(\text{fix}|a) \approx \int_{x_c}^{1/2} P_s(\text{est}|x_0, a) \cdot \rho_{msdb}(x_0|a) dx_0. \quad \text{(S16)}$$

Our derivations thus far clarify that for a segregating allele to fix with appreciable probability, it has to establish with appreciable probability, *i.e.*, $x_{msdb} \gtrsim x_{est}$ and $x_{est} > 0$ (or equivalently, if $\Lambda \gtrsim a$), and then it has to continue to fixation after it establishes, *i.e.*, $x_{msdb} \gtrsim x_c$.

We can derive a rough yet insightful analytic approximation for the critical frequency $x_c$. To this end, we approximate the change in allele frequency deterministically in continuous time, assuming that it driven by directional selection alone. Namely,

$$\frac{dx}{dt} = \frac{a \cdot D}{V_S} x(1-x). \quad \text{(S17)}$$

In turn, the change in the distance of the population's phenotypic mean from the optimum becomes

$$\frac{dD}{dt} = -\frac{\sigma^2}{V_S} D - 2a\frac{dx}{dt} = -\left(\sigma^2 + 2a^2 x(1-x)\right) \cdot D/V_S. \quad \text{(S18)}$$

We divide Eq. S18 by Eq. S17 to obtain a simple differential equation:

$$\frac{dD}{dx} = -\left(\frac{\sigma^2/a}{x(1-x)} + 2a\right). \quad \text{(S19)}$$

We require that at the time of the shift, the allele is at the critical frequency and the distance from the optimum is the shift size, *i.e.*,

$$x(0) = x_c \text{ and } D(0) \equiv \Lambda, \quad \text{(S20)}$$

and that at the time the allele reaches frequency ½, the distance to the optimum hits 0, *i.e.*,

$$x(t_{1/2}) \equiv 1/2 \text{ and } D(t_{1/2}) = 0. \quad \text{(S21)}$$

We solve Eq. S19 to obtain an implicit equation relating the critical initial frequency to the background variance, shift size and allele effect size:

$$\Lambda \approx \text{Ln}\left(\frac{1-x_c}{x_c}\right) \cdot \frac{\sigma^2}{a} + a(1-2x_c). \quad \text{(S22)}$$

When $x_c \ll 1$, which is necessary for an appreciable probability of fixation, Eq. S22 simplifies to

$$\Lambda \approx -\text{Ln}(x_c) \cdot \sigma^2/a + a, \quad \text{(S23)}$$

and the critical frequency is approximated by

$$x_c \approx \text{Exp}(-a(\Lambda - a)/\sigma^2). \quad \text{(S24)}$$

Substituting this approximation for $x_c$ into Eq. S16 for the probability of fixation, we find that

$$P_s(\mathrm{fix}|a) \approx C\left(h\left(\frac{x_c}{x_{msdb}}\right) - h\left(\frac{x_c}{x_{msdb}} + \frac{x_c}{x_{est}}\right) + h\left(\frac{1}{2x_{msdb}} + \frac{1}{2x_{est}}\right) - h\left(\frac{1}{2x_{msdb}}\right)\right), \qquad \text{(S25)}$$

where, again, the last two terms are negligible, such that

$$P_s(\mathrm{fix}|a) \approx C\left(h\left(\frac{x_c}{x_{msdb}}\right) - h\left(\frac{x_c}{x_{msdb}} + \frac{x_c}{x_{est}}\right)\right). \qquad \text{(S26)}$$

As we would expect, the probability of fixation becomes negligible when $x_c \gg x_{msdb}$. If instead $x_c \ll x_{est}$ and $x_{msdb}$, we can approximate the probability of fixation using a first-order expansion around $x_c = 0$, such that

$$P_s(\mathrm{fix}|a) \approx C\left(\mathrm{Ln}\left(1 + \frac{x_{msdb}}{x_{est}}\right) - \frac{x_c}{x_{est}}\right). \qquad \text{(S27)}$$

When $x_{est}$ and $x_{msdb}$ are of a similar order as $x_c$, a more intricate approach is required. We can try a linear expansion around any point $x_c/x_{msdb} \le x^* \le x_c/x_{msdb} + x_c/x_{est}$. Setting $x^* = x_c/x_{msdb} + \zeta \cdot x_c/x_e$, yields

$$P_s(\mathrm{fix}|a) \approx \frac{C}{\zeta + \frac{x_{est}}{x_{msdb}}} \cdot \mathrm{Exp}\left(-\frac{x_c}{x_{est}}\left(f + \frac{x_{est}}{x_{msdb}}\right)\right). \qquad \text{(S28)}$$

A cursory analysis suggests that setting $\zeta = 1/3$ works fairly well when $\sigma^2 < a^2$.

We obtain a more accurate approximation for $x_c$ using a double recursion of the deterministic allelic and phenotypic equations (Eqs. 1 and S6). That is, we deterministically evolve the distance and the allele's frequency. Then, we use a numerical line search to find the value of $x_c$, where starting the recursions with $x_0 = x_c$ and $D(0) = \Lambda$ results in $x(t_{1/2}) = 1/2$ and $D(t_{1/2}) = 0$ (within a specified tolerance). This approach accounts for the effects of stabilizing selection on the trajectory of the allele.

In Fig. S4, we compare these analytical approximations for the fixation probability with the double recursion approximation, which accurately describes simulation results (Fig. 3D and E). Because our rough approximation of $x_c$ ignores the effects of stabilizing selection, it results in an overestimation of the fixation probability. We find this approximation insightful nonetheless, given its simplicity and that for a large effect allele to fix, directional selection must be much stronger than stabilizing selection at the time of the shift.

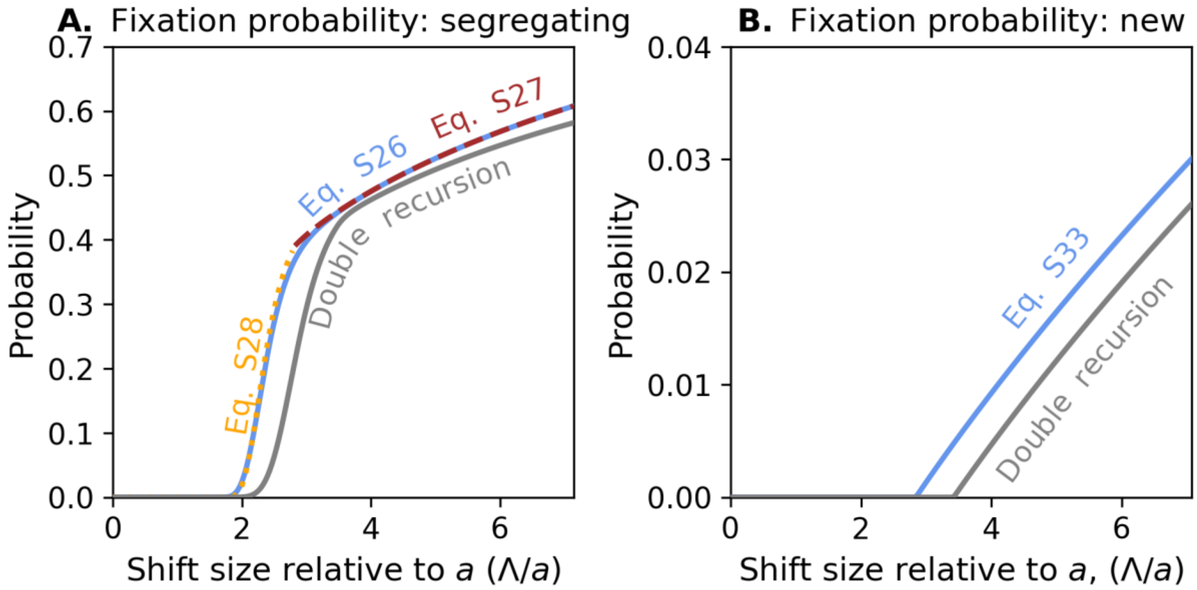

**Figure S4.** Comparison of approximations of the fixation probability for (A) segregating and (B) new alleles. These approximations assumed $a^2 = 100, N = 5000,$ and $\sigma^2 = 40$.

**The effects of drift on fixation.** The numeric approximation is quite accurate despite ignoring the effects of drift other than on the probability of establishment. One case in which we might expect drift to have a substantial effect is when the allele is initially sufficiently rare, *i.e.*, when $x_0 \lesssim 5/(2Ns_0)$ . In this case, we expect that conditional on establishment drift would introduce substantial stochasticity in its initial rate of ascent, which could affect whether it reaches frequency ½ before directional selection on it becomes negligible. We expect this to have little effect on the accuracy of our approximation because alleles that start at sufficiently low frequencies to be affected contribute negligibly to the overall probability of fixation.

The other case in which we might expect drift to substantially affect fixation is when selection on the allele becomes sufficiently weak (*i.e.*, when $2Ns \lesssim 1$) when it nears frequency ½. Notably, if drift dominates over selection at frequency ½, then we would actually expect the allele to fix with a probability of ~½. However, we would expect the probability that the allele starts at a frequency that lands it near frequency ½ when selection is sufficiently weak to be small. Furthermore, we would expect alleles that meet this condition to be nearly evenly split between those that fall slightly above and below frequency ½, such that the net effect on the probability of fixation is well approximated by our assumption that all those above frequency ½ fix and those below it are lost.

#### 5.2. The fixation probability of a new mutation

A large effect allele that arises from a mutation after the shift in optimum always starts at frequency $1/2N$, but the time at which it arises crucially affects its probability of fixing. We calculate this probability assuming that this time is uniformly distributed within the time-window in which the distance to the new optimum in the absence of large effect alleles is sufficiently large, specifically that $D_L(t) \geq \delta$. Namely, that $t_{max} \geq t \geq 0$, with $t_{max} = (V_S/\sigma^2) \cdot \mathrm{Ln}(\Lambda)$. Using a different time-window, with $t'_{max}$, should adjust the probability by a factor of (approximately) $t'_{max}/t_{max}$ so long as the probability of fixation when $t > t'_{max}$ is vanishingly small. Our derivations are generally similar to those we used for a segregating allele.

**The probability of establishment.** Similar to the case of a segregating allele, we approximate the probability of establishment for an allele that arises at time $t$ using Kimura's formula for the probability of fixation (Eq. S7), where in this case the initial selection coefficient is

$$s_0 \approx (a/V_S) \cdot D_L(t) - (a^2/V_S)(1/2 - 1/2N) \tag{S29}$$

and the population-scaled $s_0$ is approximately

$$2Ns_0 \approx a(D_L(t) - a/2), \tag{S30}$$

with $D_L(t) = \Lambda \cdot \mathrm{Exp}(-(\sigma^2/V_S)t)$. The probability of establishment is therefore well approximated by

$$P_n(\mathrm{est}|t,a) \approx 1 - \mathrm{Exp}\big(-2(a/V_S)(\Lambda \cdot \mathrm{Exp}(-(\sigma^2/V_S)t) - a/2)\big), \tag{S31}$$

For a new allele to establish with appreciable probability, the selection favoring it must be strong relative to genetic drift, *i.e.*, $2Ns_0 \approx a(D_L(t) - a/2) \gg 1$. No new allele can establish when the shift size $D_L(0) = \Lambda < a/2$, and for a new allele to establish it must arise before

$$t_{est} = (V_S/\sigma^2) \cdot \mathrm{Ln}(2\Lambda/a). \tag{S32}$$

For a new allele to establish with appreciable probability, it must arise when $t < (V_S/\sigma^2) \cdot \mathrm{Ln}(\Lambda/(10/a + a/2))$.

We approximate the probability that a new allele establishes—without conditioning on the time it arises—by integrating over the times it can arise. Assuming that $t_{max} \geq t_{est} > 0$ (otherwise none would establish), this probability is

$$P_n(est|a) \approx \frac{1}{t_{max}} \int_0^{t_{est}} P_n(est|t,a) \cdot dt \tag{S33}$$

$$= \frac{\mathrm{Ln}\left(\frac{2\Lambda}{a}\right)}{\mathrm{Ln}(\Lambda)}\left(\mathrm{Ln}\left(\frac{2\Lambda}{a}\right) - e^{a^2/V_S}\left(h\left(\frac{a^2}{V_S}\right) - h\left(\frac{2a\Lambda}{V_S}\right)\right)\right).$$

The probability of establishment does not depend on the background variance, $\sigma^2$, because per our definition of $t_{max}$, its effects on $t_{est}$ and $t_{max}$ cancel out.

**The probability of reaching frequency ½.** By analogy with the case of a segregating allele, if we neglect the stochasticity in the allele's rate of ascent at low frequencies (*e.g.*, during establishment), we expect there to be some critical time, $t_c$, such that an allele that arises at this time and establishes will reach frequency ½ when the distance to the optimum hits 0. Alleles that arise before this critical time will eventually fix, whereas those that arise after it will be lost. We can therefore approximate the probability that a new allele with effect size $a$ fixes by

$$P_n(\mathrm{fix}|a) \approx \frac{1}{t_{max}} \int_0^{t_c} P_n(\mathrm{est}|t, a)dt. \tag{S34}$$

Following the rough analytic derivation we used to approximate $x_c$ for a segregating allele but replacing the initial conditions with shift size $D_L(t_c) = \Lambda \cdot \mathrm{Exp}(-(\sigma^2/V_S)t_c)$ and frequency $x(t_c) = 1/2N$, we find that

$$t_c \approx (V_S/\sigma^2) \cdot \mathrm{Ln}\left(\frac{\Lambda}{\mathrm{Ln}(2N)\cdot\sigma^2/a + a}\right). \tag{S35}$$

This approximation suggests that a new allele can fix only if $\Lambda > \mathrm{Ln}\,(2N) \cdot \sigma^2/a + a$, such that $t_c > 0$. Substiting this approximation for $t_c$ into Eq. S34 for the probability of fixation, we find that

$$P_n(\mathrm{fix}|a) \approx \frac{1}{\mathrm{Ln}(\Lambda)}\left(\mathrm{Ln}\left(\frac{\Lambda}{\mathrm{Ln}(2N)\cdot\sigma^2/a + a}\right) - e^{a^2/V_S}\left(h\left(\frac{a^2}{V_S}\right) - h\left(\frac{2\left(\mathrm{Ln}(2N)\cdot\sigma^2 + a^2\right)}{V_S}\right)\right)\right). \tag{S36}$$

Similar to the way we obtained a more accurate of $x_c$, here we use a numeric approach to obtain a more accurate approximation of $t_c$. In this case, however, accounting for the accelerated ascent of an allele that establishes when it is rare substantially improves the accuracy of our approximation, because new alleles, which start at frequency $1/2N$, are more substantially affected by genetic drift than segregating alleles. We account for this acceleration by dividing the trajectory of alleles that establish into a 'stochastic' phase up until frequency $1/s_0$ and a 'deterministic' phase above this frequency.

We calculate the expected time for a new mutation to reach frequency $1/s_0$ using the diffusion approximation (Chapter 4 in Ewens 2004). To this end, we treat $x_{est} = 1/s_0$ and $x = 0$ as absorbing boundaries, and approximate the expected time for reaching frequency $x_{est}$ as

$$\bar{\tau}(x_0|x_\infty = x_{est}) = \int_0^{x_{est}} \tau(x|x_0 = 1/2N\,, x_\infty = x_{est})dx, \quad \text{(S37)}$$

where $\tau(x|x_0 = 1/2N\,, x_\infty = x_{est})dx$ is the expected sojourn time that an allele spends between frequencies $x$ and $x + dx$ conditional on eventually reaching frequency $x_{est}$. The conditional sojourn time density can be expressed as

$$\tau(x|x_0 = 1/2N\,, x_\infty = x_{est}) = \tau^*(x|x_0)\frac{P_{x_{est}}(x)}{P_{x_{est}}(x_0)}, \quad \text{(S38)}$$

where $\tau^*(x|x_0)$ is the density of sojourn time conditional on reaching *either boundary* and $P_{x_{est}}(x)$ is the probability of an allele starting at frequency $x$ reaching frequency $x_{est}$. We approximate $\tau^*(x|x_0)$ and $P_{x_{est}}(x)$ based on the first two moments of change in allele frequency per generation (Eqs. 1 and 2) assuming a constant distance to the new optimum during this phase. Specifically, we substitute $D = D_L(t_0) = \Lambda \cdot \mathrm{Exp}(-(\sigma^2/V_S)t_0)$ into $E(\Delta x|x)$, which makes both moments independent of time. In this approximation,

$$\tau^*(x|x_0) \approx \begin{cases} \dfrac{2P_0(x_0)}{E(\Delta x|x)\psi(x)}\displaystyle\int_0^x \psi(y)dy\,, & 0 \le x \le x_0 \\ \dfrac{2P_{x_{est}}(x_0)}{E(\Delta x|x)\psi(x)}\displaystyle\int_x^{x_{est}} \psi(y)dy\,, & x_0 \le x \le x_{est} \end{cases} \quad \text{(S39)}$$

and

$$P_{x_{est}}(x) = \frac{\int_0^x \psi(y)dy}{\int_0^{x_{est}} \psi(y)dy}, \quad \text{(S40)}$$

where $\psi(x) = \mathrm{Exp}\left(-2\int_0^x \frac{E(\Delta x|x)}{V(\Delta x|x)}dx\right)$ and $P_0(x) = 1 - P_{x_{est}}(x)$. We perform these integrals numerically to obtain $\bar{\tau}(x_0|x_\infty = x_{est})$, given the shift size and the allele's effect size and time of appearance.

We approximate the remainder of the allele's trajectory using the double recursion of the deterministic allelic and phenotypic equations (Eqs. 1 and S6). We start the recursion with the initial conditions after establishment, *i.e.*, with $x(0) = 1/s_0$ and $D(0) = \Lambda \cdot \mathrm{Exp}\left(-(\sigma^2/V_S)\left(t_0 + \bar{\tau}(x_0|x_\infty = x_{est})\right)\right) - 2a(1/s_0 - 1/2N)$, and use it to ascertain whether the allele reaches frequency ½ and to calculate $D\left(t_{1/2}\right)$ when it does (with $t_{1/2}$ defined by $x\left(t_{1/2}\right) = 1/2$). We perform a line search on $t_0$ to approximate the critical time $t_0 = t_c$ such that $D\left(t_{1/2}\right) = 0$.

Figure S6 shows a comparison of our approximations for the fixation probability of a new allele with simulations. As we would expect our rough analytic approximation overestimates the fixation probability because it neglects the effects of stabilizing selection. In turn, our numerical approximation performs quite well.

#### 5.3. Allele trajectories

As we already noted, a large effect allele can take different kinds of trajectories toward its ultimate fixation or loss. Here, we focus on segregating alleles, noting that the derivation for a new allele arising at time $t_0$ would be the same except for replacing $D(0) = \Lambda$ with $D(0) = D_L(t_0)$.

The classification into different kinds of trajectories and the conditions for each are easiest to understand when the background variance $\sigma^2$ is tiny (Fig. S5A and B). In this case, the background's initial contribution to adaptation is negligible, which allows us to approximate the population's mean distance to the optimum by $D \approx \Lambda - 2a(x - x_0)$ and the allele's change in frequency per generation by

$$E(\Delta x|x) \approx a/V_S \cdot \Big(\big(\Lambda - 2a(x - x_0)\big) - a(1/2 - x)\Big)\, x(1 - x). \quad \text{(S41)}$$

This expression is null, with the effects of stabilizing and directional selection on allele frequency canceling out, when

$$x^* \approx \Lambda/a - 1/2 + 2x_0. \quad \text{(S42)}$$

When $x^* < 0$ the allele can never establish and is destined to *quick loss*. In all other cases, the allele can establish with some probability, but failing to do so would also lead to a quick loss. Henceforth, we consider only alleles that establish.

Next, consider the cases in which $1 > x^* > 0$, or, equivalently, when $3a/2 > \Lambda > a/2$ (where, for simplicity, we drop $x_0$ here and below, given that $x_0 \ll 1$). Here, frequency $x^*$ is a stable fixed point, because $E(\Delta x|x^*) = 0$ and $E'(\Delta x|x^*) < 0$. Once the allele reaches it, its frequency will be held near $x^*$ by balancing selection, with some fluctuations due to genetic drift. At the lower half of this range, when the shift size is $a > \Lambda > a/2$, the allele reaches the fixed point at $0 < x^* < 1/2$ before the population mean reaches the new optimum. Adaptation is then mediated by background alone, which pushes the population mean to the new optimum over a timescale of $V_S/\sigma^2$ generations. As the distance from the optimum decreases, directional selection on the allele weakens, which reduces $x^*$. This process continues until the allele is lost ($x^* \approx 0$ when we substitute $D \approx a/2$ for $\Lambda$ in Eq. S42). In this case, the allele is established and then lost.

At the higher half of this range, when the shift size $3a/2 > \Lambda > a$, the allele reaches the fixed point at $1/2 < x^* < 1$ when the population mean has already overshot the optimum. In this case, the background pushes the population mean back to the new optimum over a timescale of $V_S/\sigma^2$. As the distance from the optimum decreases, directional selection on the allele, which is now negative, weakens, and stabilizing selection acts to increase the frequency $x^*$, until the allele fixes ($x^* \approx 1$ when we substitute $D \approx 3a/2$ for $\Lambda$ in Eq. S42). In this case, an allele experiences a partial selective sweep.

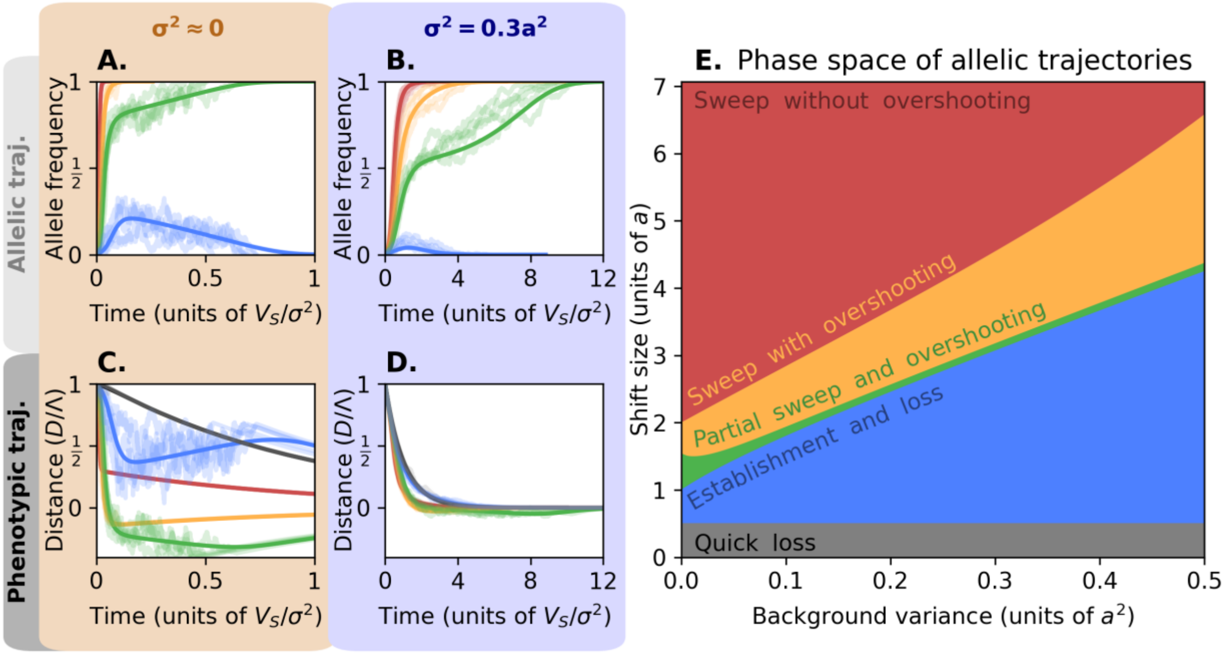

**Figure S5. Allelic and phenotypic trajectories**. We illustrate the different kinds of allelic and phenotypic trajectories for a large effect allele, with $a^2 = 200\delta^2$ and initial frequency $x_0 = 1/a^2$), with tiny background variance $\sigma^2 = 1\delta^2$ (A and B), and with appreciable background variance, $\sigma^2 = 30\delta^2 = 0.3a^2/2$ (C and D). The different kinds of trajectories in each case correspond to different shift sizes, with a constant population size of $N = 5000$. We show both the expected trajectories, which are calculated using a double recursion (Eqs. 1 and S6), and simulated trajectories, which were generated using *single allele* simulations. The trajectories of alleles that experience quick loss would not be visible and are therefore not plotted. (E) A phase space of adaptive trajectories. The combinations of shift sizes and background variances resulting in different kinds of trajectories are calculated based on the double recursion, with other parameter values similar to those in A-D.

When $x^* > 1$, or, equivalently, when $\Lambda > 3a/2$, the allele is subject to positive selection all the way to fixation; it experiences a full selective sweep. We split this case further based on the phenotypic trajectories. When the shift size is $3a/2 < \Lambda < 2a$, the population mean overshoots the optimum, and when $\Lambda > 2a$ it does not overshoot.

All of these trajectories also occur when the background variance is appreciable (Figs. S5B, D, and E). As we already saw, the minimal shift size that allows the allele to establish is insensitive to the background variance, and the minimal shift size that allows it to fix increases approximately linearly with the background variance (Eq. S23). Similar reasoning indicates that the minimal shift size that allows for a full sweep with and without overshooting the optimum also increases approximately linearly with the background variance. Lastly, when the background variance increases, more rapid adaptation from the background narrows down the range of shift sizes in which the allele experiences a partial sweep with the phenotypic mean overshooting the optimum.

The transient balancing selection we described is a special case of adaptive heterozygote advantage previously described by Sellis et al. (2011). The partial and full sweeps we described should affect levels of linked neutral diversity along the lines describe by Barton (2000) and Coop and Ralph (2012).

### 6. The general case of large effect alleles and a Fisherian background

Figure S6 shows how the mutational input of large effect alleles affects the probability that any large effect allele reaches fixation and their long-term (fixed) adaptive contribution for three combinations of shift sizes and background variances. As we would expect, this contribution increases when the shift size increases or the background variance decreases. In all cases and as we already saw for one of them in Fig. 4A, when the mutational input is low (in the shaded region corresponding to $2NU < 1$) the adaptive contribution increases with the mutational input. In section 6.1, we derive a simple approximation for the number of segregating and new large effect alleles that fix in this parameter range and their adaptive contributions (dashed lines in panels of Fig. S6); this is also the approximation that we used in Figs. 4B and C.

In all cases, when the mutational input increases further (*i.e.*, when $2NU \gtrsim 1$) the long-term adaptive contribution of large effect alleles decreases with increasing mutational input. This decline reflects interference among multiple large effect alleles, which is mediated through their effects on the phenotypic dynamic. In section 6.2, we consider the phenotypic dynamic and its dependence on model parameters.

The mutational input also affects the characteristics of the alleles that fix. Notably, in parts of the range where $2NU \gtrsim 1$, the number of large effect fixations may increase while their adaptive contribution decreases with increasing mutational input (compare Figs. S6A and D), indicating changes to the distribution of effect sizes of the alleles that fix. Additionally, the mutational input also affects how fixations are distributed among segregating and newly arising large effect alleles as seen in Figs. S6B and C. In section 6.3, we consider how model parameters affect the characteristics of fixations.

Lastly, some allele effect sizes are too large for their effects to be well approximated by the Fisherian background but too small to be considered large. In section 6.4, we consider how such alleles affect the fixation of large effect alleles, as well as their fixation and adaptive contribution.

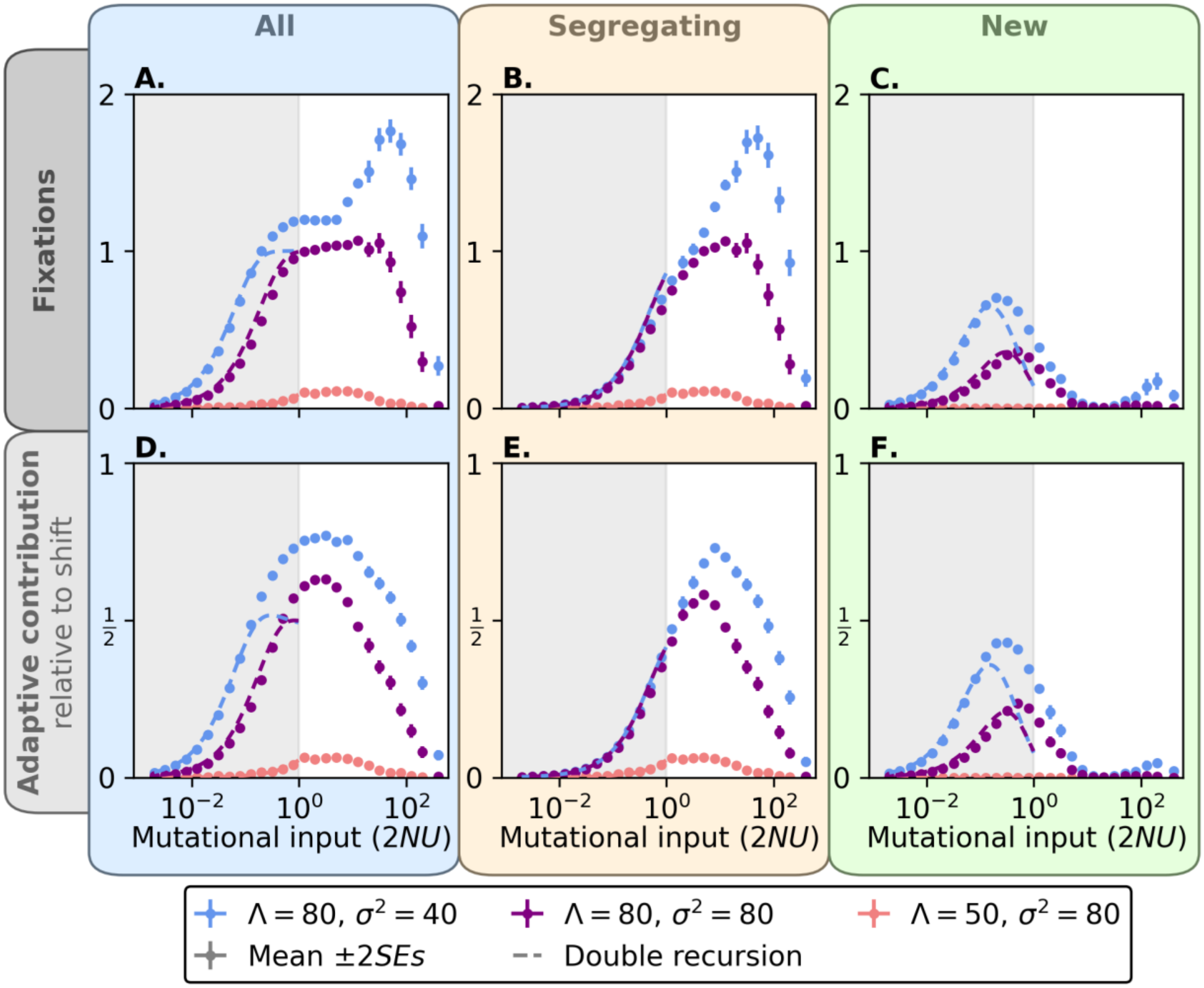

**Figure S6. Long-term adaptive response of large effect alleles vs. their mutational input.** The number of fixations (top) and their contribution to adaptation relative to the shift size (bottom) for all (left), segregating (middle) and new (right) large effect alleles, for three combinations of shift sizes and background variances. Simulation results show the mean ± 2SEs (unless they SEs are too small) calculated over ≥ 600 replicates (and up to 10000 for 2NU ≪ 1) for each combination of parameters. The analytic approximations for $2NU < 1$ (dashed curves) were calculated using Eqs. S46 and S49 for the number of fixations; similar integrals but including a $2a(1 - x_0)$ term were numerically evaluated to calculate the expected adaptive contribution.

### 6.1. Low mutational input ($2NU < 1$)

At MSDB, before the shift, large effect alleles follow stochastic frequency trajectories, starting from a new mutation and ending with loss. The sojourn time of an allele with effect size $a$ at frequency $x$, $\tau_{msdb}(x|a)$, is defined such that $\tau_{msdb}(x|a) \cdot dx$ is the expected time that the allele spends between frequencies $x$ and $x + dx$ throughout its trajectory (chapter 4, Ewens 2004). We denote the expected total time an allele spends in the population under MSDB by $\bar{\tau}_{msdb}(a)$. The sojourn time relates to the frequency density, which we used in section 5.1, by

$$\tau_{msdb}(x|a) = \bar{\tau}_{msdb}(a) \cdot \rho_{msdb}(x|a) \quad \text{(S43)}$$

and is approximated in Appendix 3 of Hayward and Sella (2022). We use $\bar{\tau}_{msdb}(a)$ to relate the expected number of fixations in the low mutational input range with our approximations for a single large effect allele (section 5.1).

When the mutational input is sufficiently low, multiple segregating and new large effect alleles rarely interfere with one another's chance of fixing after the shift in optimum. If we ignore interference, we can approximate the expected number of segregating alleles that establish as

$$E(n_{e|s}) \approx 2NU \cdot \int_{a_{min}}^{\infty} P_s(\text{est}|a)\, \bar{\tau}_{msdb}(a)\, g(a) da. \quad \text{(S44)}$$

We approximated $P_s(\text{est}|a)$—the probability that a single segregating allele with effect size $a$ establishes—in section 5.1 (Eq. S15), and the factor $2NU \cdot \bar{\tau}_{msdb}(a)$ is the expected number of alleles with effect size $a$ segregating before the shift. The lower limit of the integration $a_{min}$ denotes the minimal effect size that we include in the large effect mode of the model; in most simulations we take $a_{min} = 10$ but we consider lower limits below (section 6.4).

By the same token, we can approximate the expected density of segregating alleles with effect size $a$ that fix as

$$E(n_{f|s}|a) \approx 2NU \cdot \bar{\tau}(a) \cdot P_s(\text{fix}|a), \quad \text{(S45)}$$

and the expected total number of segregating alleles that fix as

$$E(n_{f|s}) \approx 2NU \cdot \int_{a_{min}}^{\infty} P_s(\text{fix}|a)\, \bar{\tau}(a)\, g(a) da, \quad \text{(S46)}$$

where we approximated $P_s(\text{fix}|a)$—the probability that a single segregating allele with effect size $a$ fixes—in section 5.1 (Eq. S16).

We can derive similar approximations for newly arising alleles. We approximate the expected number of new alleles that establish by

$$E\left(n_{e|n}\right) = 2NU \cdot t_{max} \cdot \int_{a_{min}}^{\infty} P_n(\text{est}|a)\, g(a) da, \tag{S47}$$

where we approximated $P_n(\text{est}|a)$—the probability that a new mutation with effect size $a$ establishes—in section 5.2 (Eq. S31), and the factor $2NU \cdot t_{max}$ is the expected number of large effect mutations that arise within $t_{max}$ generations after the shift. Similarly, we approximate the expected density of new alleles with effect size $a$ that fix as

$$E\left(n_{f|n}|a\right) \approx 2NU \cdot t_{max} \cdot P_n(\text{fix}|a), \tag{S48}$$

and the expected total number of new alleles that fix as

$$E\left(n_{f|n}\right) = 2NU \cdot t_{max} \cdot \int_{a_{ml}}^{\infty} P_n(\text{fix}|a)\, g(a) da, \tag{S49}$$

where we approximated $P_n(\text{fix}|a)$—the probability that a new mutation with effect size $a$ fixes—in section 5.2 (Eq. S34).

When the mutational input is sufficiently low for these approximations to apply, the expected numbers of fixations from both segregating and new mutations are proportional to the mutational input $2NU$, where the proportionality constant depends on the shift size and background variance, and the number of segregating and new mutations to establish and fix follow Poisson distributions with the expectations we calculated.

Once one large effect allele reaches fixation, however, the directional selection that future large effect alleles experience is weakened. To extend the scope of our approximation to somewhat higher mutational inputs, we account for this kind of interference by assuming that only one large effect allele can fix. Specifically, we assume that if the number of alleles that would have fixed independently is greater than one then exactly one fixes, and if a segregating allele fixes, new mutations do not. In this approximation, the expected number of fixations equals the probability of fixation. The expected number of segregating fixations equals the probability that not all segregating alleles go extinct (assuming independence), *i.e.*,

$$P\left(n_{f|s} = 1\right) = 1 - \text{Exp}\left(-E\left(n_{f|s}\right)\right). \tag{S50}$$

The expected number of new mutations to fix equals the probability that all segregating alleles—but not all new alleles—go extinct (again assuming independence), which is

$$P\left(n_{f|n} = 1\right) = \underbrace{\text{Exp}\left(-E\left(n_{f|s}\right)\right)}_{\substack{\text{Probability that all} \\ \text{segregating alleles} \\ \text{go extinct}}} \cdot \underbrace{\left(1 - \text{Exp}\left(-E\left(n_{f|n}\right)\right)\right)}_{\substack{\text{Probability at least} \\ \text{one new allele} \\ \text{reaches fixation}}}. \tag{S51}$$

The expected total number of fixations is the sum of these two expectations

(S52)

$$P(n_f = 1) = P(n_{f|s} = 1) + P(n_{f|n} = 1) = 1 - \mathrm{Exp}\left(-\left(E(n_{f|s}) + E(n_{f|n})\right)\right).$$

In Fig. 4B and C, we use these approximations to show how the expected number of fixed and established large effect alleles depends on the shift size, and in Fig. S7 we show that for a mutational input of $2NU = 0.01$ used in Fig. 4, these approximations are in good agreement with simulations.

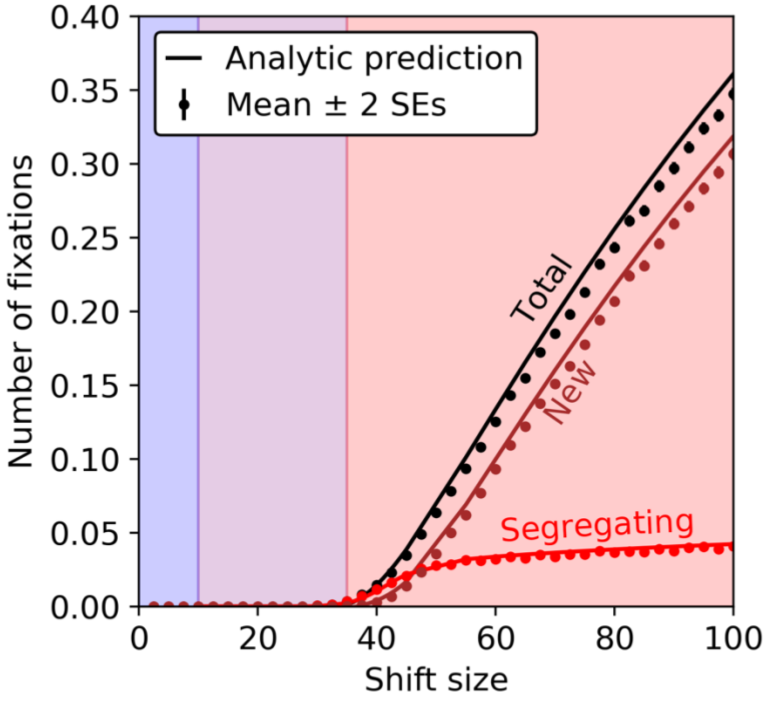

**Figure S7. Number of alleles that fix with low mutational input.** This figure is the same as Fig. 4B, other than also showing the mean number of fixations in simulations. The simulations are as described in Fig. 4.

These approximations capture the qualitative dependence of the number of fixations on the mutational input when $2NU \lesssim 1$ (dashed lines in Figs. S6A-C). They show that the number of fixations from segregating alleles increases with mutational input throughout this range. They also show that when the mutational input is tiny, the number of fixations from new alleles increases with mutational input so long as $t_c > 0$ (otherwise, new mutations cannot fix). Lastly, they show that once the mutational input is large enough for fixations from segregating alleles to become common, these fixations cause a reduction in the number of fixations from new alleles.

Our approximations are quantitatively accurate when $2NU < 0.1$ but their accuracy diminishes when $1 > 2NU > 0.1$. In the latter range, multiple large effect alleles can fix and/or new alleles can fix before segregating alleles, thus violating the assumptions of our approximation. Consequently, we underestimate the number of fixations from new alleles and in some cases the number of fixations from segregating alleles. At larger mutational inputs ($2NU \gtrsim 1$), multiple segregating large effect alleles can establish and cause substantial interference, which alters the probability in fixations in ways that the approximations of this section do not describe.

#### 6.2. High mutational input ($2NU > 1$)

As we illustrated in the main text, when the mutational input is large, the adaptive response involves substantial changes to the shape of the phenotypic distribution, notably to the phenotypic variance and skew. Before the shift, at MSDB, the expected phenotypic variance

$$V_A(0) = 2NU \cdot \int_{a_{ml}}^{\infty} \left( \int_0^{1/2} v(a,x) \cdot \tau_{msdb}(a,x) \cdot dx \right) g(a) \cdot da + \sigma^2, \quad \text{(S53)}$$

where $v(a,x) = 2a^2x(1-x)$ is the contribution to variance of a variant with effect size $a$ at frequency $x$. The expected contribution of large effect variants to phenotypic variance per unit mutational input is approximately constant, with

$$\int_0^{1/2} v(a,x) \cdot \tau_{msdb}(a,x) \cdot dx \approx 4 \quad \text{(S54)}$$

(see, *e.g.*, Fig. 3 in Simons et al. (2018)) and therefore

$$V_A(0) \approx 8NU + \sigma^2. \quad \text{(S55)}$$

The phenotypic skew has an expectation of zero, with contributions of aligned and opposing alleles canceling each other out (Appendix 3 in Hayward and Sella (2022)).

After the shift, directional selection increases the frequency of large effect alleles that are aligned with shift relative to alleles with opposing effects. This manifests in individuals carrying more large effect alleles on average, with greater variability in their number among individuals, which dramatically increases the phenotypic variance. Individuals also carry more alleles with aligned effects than opposing ones, which dramatically increases the phenotypic skew. When the population mean phenotype is sufficiently far from the optimum, the increase in phenotypic variance accelerates the approach of the mean to the new optimum whereas the increase in the phenotypic skew has little effect. However, when the population mean gets closer to the new optimum, adaptation gradually slows down, because the number of individuals with too many aligned alleles—that overshoot the optimum—gradually increases. When selection against individuals that overshoot the optimum nearly perfectly balances selection against those below it (Fig. 5B), the change in distance to the optimum (Eq. S4) nearly grinds to a halt, as

$$E(\Delta D) \approx -\frac{V_A(t)D(t)}{V_S} + \frac{\mu_3(t)}{2V_S} \approx 0. \tag{S56}$$

Hayward and Sella (2022) explore this 'quasi-static phase' in their Appendix 3.

Figs. S8 shows how our model's parameters affect the time, $t_{qs}$, and distance from the optimum, $D_{qs}$, at which the population enters the quasi-static phase. This phase begins when

$$D(t) \approx \frac{\mu_3(t)}{2V_A(t)} \tag{S57}$$

for the first time. In simulations, we average the first three central moments of the phenotypic distribution in sliding windows of 10 generations, and define the beginning of the quasistatic phase as the first time in which $|(\overline{D}(t) - \overline{\mu_3}(t)/2\overline{V_A}(t))/\overline{D}(t)| < 0.05$. Increasing the mutational input decreases $t_{qs}$, because having more large effect alleles speeds up the initial rate of adaptation and the buildup of the phenotypic skew. The trade-off between these effects maximizes $D_{qs}$ at some intermediate mutational input.

For large mutational inputs $(2NU > 10)$, the distance at which the quasistatic phase begins, $D_{qs}$, determines whether large effect alleles can fix. Alleles with effect sizes $a/2 > D_{qs}$ are negatively selected throughout the quasistatic phase and therefore cannot fix unless they near frequency ½ before this phase begins. With very large mutational inputs $(2NU \gg 1)$ $t_{qs}$ is small, so all large effect alleles will enter the quasistatic phase at low frequencies. In this case, if $a_{min}/2 > D_{qs}$ then no large effect will be able to fix. If $a_{min}/2 < D_{qs}$ then alleles with sufficiently small effect sizes will remain positively selected for a while and could potentially fix. However, as the population nears the optimum, directional selection on these alleles will gradually decrease. Most of these alleles will then experience a period of selective neutrality (with directional and stabilizing selection cancelling each other out) followed by a period of negative selection where they are driven to extinction. With moderate mutational inputs $(1 \leq 2NU \leq 10)$, $t_{qs}$ is larger, such that large effect alleles may near frequency ½ before entering the quasistatic phase, allowing a few of them to fix even if $a_{min}/2 < D_{qs}$, let alone if $a_{min}/2 > D_{qs}$. At low mutational inputs $(2NU < 1)$, the quasistatic phase begins long after the shift and is not an impediment to large effect fixations. Here, the quasistatic phase can occur after the population mean phenotype has overshot the new optimum.

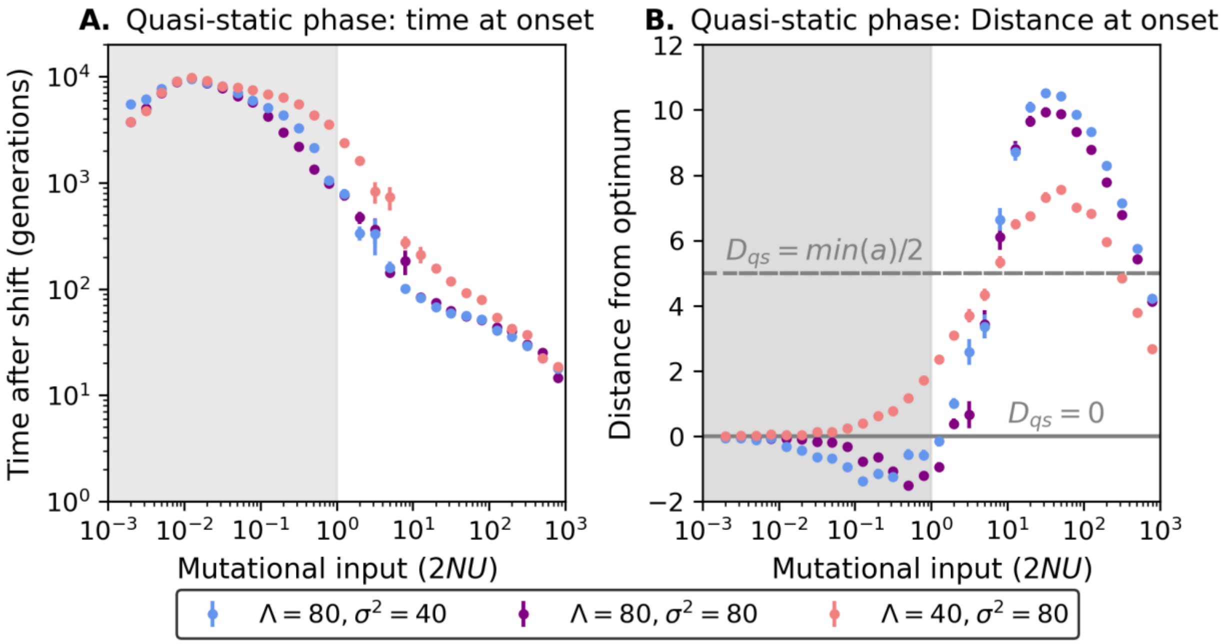

**Figs. S8. The time (A) and distance (B) when the quasistatic adaptation phase begins.** We show these summaries as functions of the mutational input for three different combinations of shift size and background variance. For the definition we use for the beginning of the quasistatic phase, see text, and for more details about the simulations, see Fig. S6. The cases in (B) in which $\overline{D}_{qs} < 0$ reflect the kind of transient overshooting we discussed supplementary section 5.3.

#### 6.3. Effect sizes of fixed alleles

We can approximation the distribution of effect sizes of large effect fixations, $g_f(a)$, in the limit of tiny mutational input, *i.e.*, when $2NU \ll 1$. The transformation between the mutational and fixed distributions of effect sizes follows from Bayes' theorem, *i.e.*,

$$g_f(a) = P(\text{fix}|a)/P(\text{fix}) \cdot g(a). \tag{S58}$$

When the mutational input is sufficiently low such that fixations are rare ($2NU \ll 1$), we assume that the fate each large effect allele is independent of the others and approximate the transformation by

$$g_f(a) = P(\text{fix}|a)/P(\text{fix}) \cdot g(a) \approx \frac{E(n_{f|s}|a)+E(n_{f|n}|a)}{\int_{a_{ml}}^{\infty}\left(E(n_{f|s}|a\prime)+E(n_{f|n}|a\prime)\right)g(a')da'}, \tag{S59}$$

where we approximated $E(n_{f|s}|a)$ and $E(n_{f|n}|a)$ in Eqs. S45 and S48, respectively. Thus, in this limit $g_f(a)$ does not depend on the mutational input. This approximation assumes no more than one fixation and breaks down when the mutational input becomes appreciable (*i.e.*, when $2NU \gtrsim 1$). Notably, it does not capture the effects of interference among large effect alleles (see below).

In this limit, the shape of the distribution of fixed effect sizes depends on the range of large effects that could potentially fix (Fig. S9). In the 'permissive' extreme, with sufficiently large shift sizes and small background variances, segregating and new alleles with any large effect size could fix. In terms of our derivations for a single large effect allele, this would be the case if $x_{msdb} \gg x_c$ and $t_c \gg 1$ for any large effect size (section 5). In this case, the transformation between $g(a)$ and $g_f(a)$ (Eq. S59) slightly favors the higher end of large effects, because new alleles account for most fixations (when $t_c \gg 1$); new alleles with larger effect sizes experience stronger directional selection and thus establish and fix with greater probabilities than those with smaller effect sizes. In the other extreme, the shift size and background variance allow only alleles in a restricted range of effect sizes to fix. In this 'restrictive' extreme, $x_c \gtrsim x_{msdb}$ and $t_c \lesssim 0$ for all effect sizes such that only alleles segregating at high frequencies at the time of the shift can fix (section 5). The transformation between $g(a)$ and $g_f(a)$ (Eq. S59) favors alleles at the lower end of large effect sizes, because they segregate at higher frequencies than alleles with larger effect sizes before the shift (and are also more abundant). As a result, the average effect size of a fixed allele is much smaller when fixation is restrictive than when it is permissive (Fig. S10).

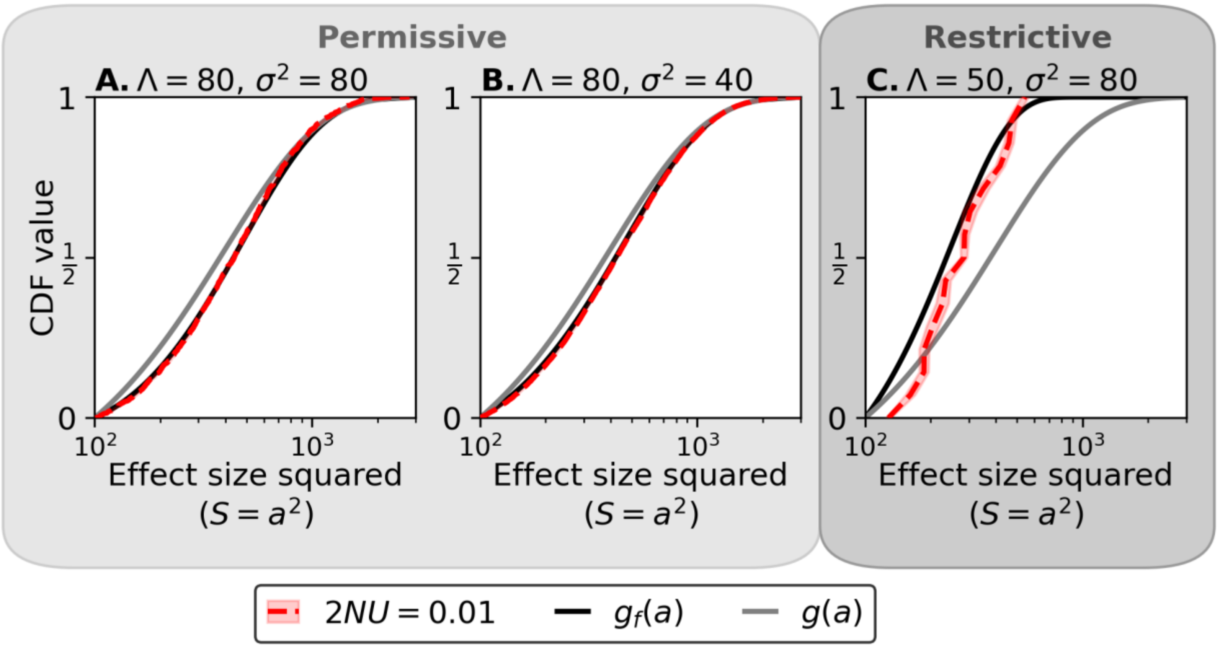

**Figure S9. Effect size distributions of fixed alleles for a tiny mutational input.** The distribution of effect sizes for fixed large effect alleles are shown for three different combinations of shift size and background variance when the mutational input of large effect alleles is sufficiently miniscule that interference should be negligible. Two of these shift size and background variance combinations correspond to the ‘permissive’ extreme (A and B) and one corresponds to the ‘restrictive’ extreme (C). Distributions are shown as CDFs. In all panels, we show the expected distribution of fixed effect sizes (Eq. S59) and the assumed distribution of effect sizes for newly arising alleles. Simulations were run as described in Fig. 4. Sleeves represent the error calculated from 10 bootstraps over the simulated data; error is larger in the ‘restrictive’ extreme (C) as far fewer alleles reach fixation.

Increasing the mutational input affects the distribution of fixed effect sizes differently in the ‘permissive’ and ‘restrictive’ extremes (Fig. S10). In the permissive extreme, when the mutational input is low ($2NU < 1$) increasing the mutational input strengthens the fixation bias toward the high end of large effects. In the restrictive extreme, when the mutational input is low ($2NU < 1$), the distribution of fixed effects is insensitive to increasing the mutational input. However, for both extremes, increasing the mutational input beyond $2NU \approx 1$ increasingly biases fixations towards alleles with smaller effect sizes (section 6.2), due to interference among large effect alleles. For very large mutational inputs (*e.g.*, $2NU = 100$), only the smallest large effect alleles can fix. These changes to the shape of distribution are reflected in the average effect size of fixed alleles.

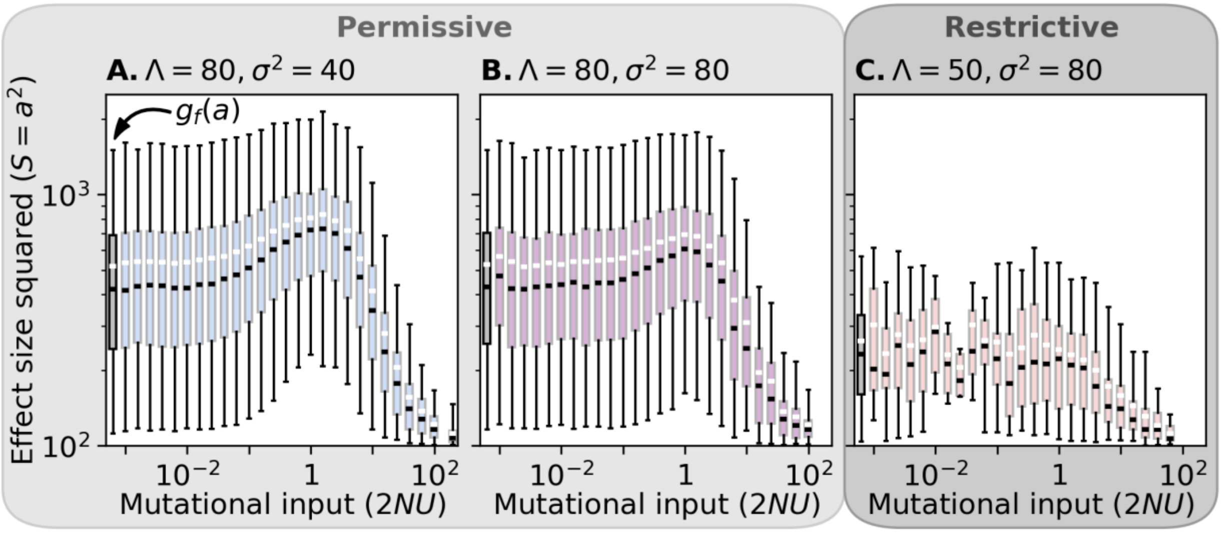

**Figure S10. Effect size distributions of fixed alleles.** The distribution of effect sizes for fixed large effect alleles are shown for three different combinations of shift size and background variance as a function of mutational input. Two of these shift size and background variance combinations correspond to the ‘permissive’ extreme (A and B) and one corresponds to the ‘restrictive’ extreme (C). Distributions are shown as boxplots where the median is indicated in black, the mean is indicated in white, the interquatile range is represent by a filled box and the whiskers represent the 95$^{th}$ interpercentile range. In all panels, we show the expected distribution of fixed effect sizes (Eq. S59) in grey at the left most edge. Simulations were run as described in Fig. 4. When the number of realized fixations is too small to estimate the components of the 95th interpercentile range, the whiskers are drawn from the minimum and maximum fixed effect size. Parameter combinations with fewer than five fixations are not shown.

#### 6.4. Midrange effect alleles

The distributions of large effect sizes we used do not include effect sizes between $a^2 = 10$ and $100$, which are also too large to be well approximated in terms of the background variance; we refer to these effects as 'midrange'. Figure S11 shows the genetic basis of the long-term adaptive response in simulations that include midrange alleles, where we divide their effect sizes into 'low midrange' with $10 < a^2 < 30$ and 'high midrange' with $30 < a^2 < 100$ for reasons that will become apparent below.

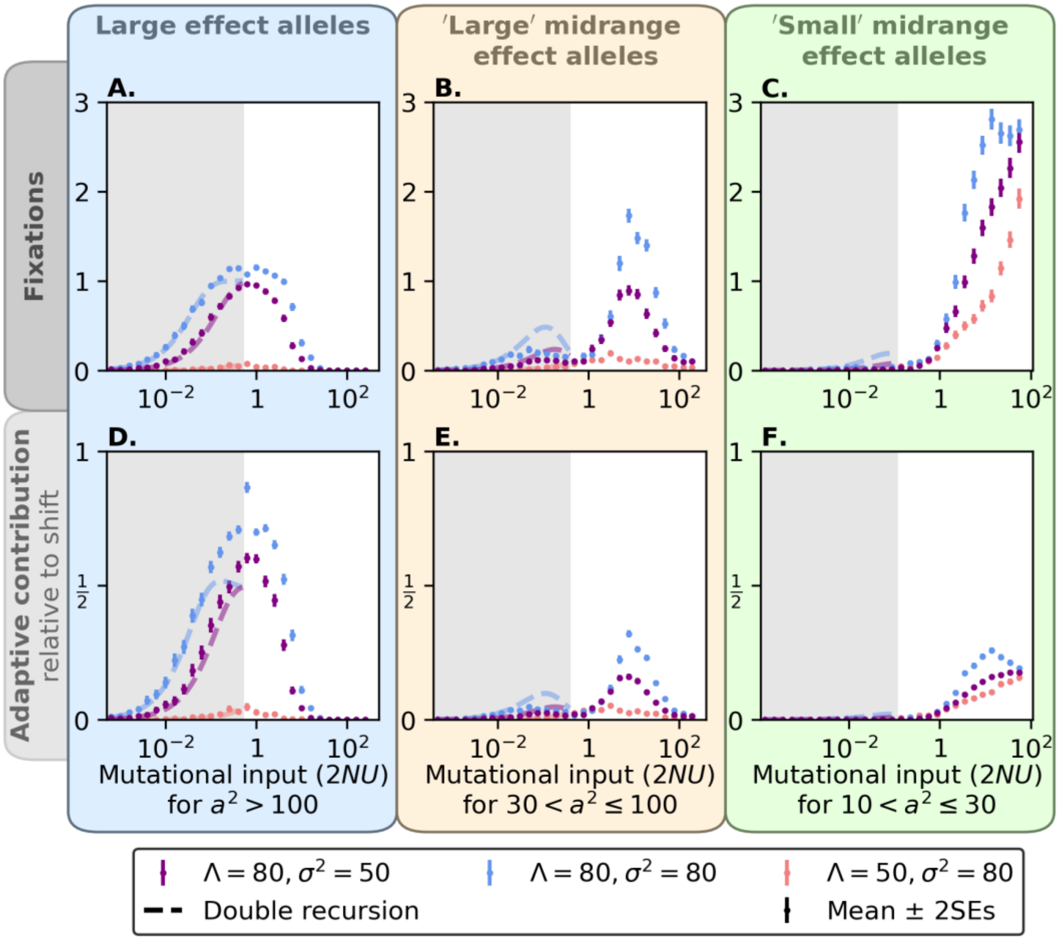

**Figure S11. The genetic basis of adaptation with midrange effect alleles**. We show the number of fixations and long-term adaptive contribution as a function of the mutational input for each class of alleles for three different combinations of shift size and background variance. The mutational distribution of effect sizes was taken to be an equal mix between the large effect distribution we use in the main text ($a^2 = 100 + \tilde{a}^2$ with $\tilde{a}^2 \sim Exp(400)$) and a uniform distribution for midrange effects ($a^2 \sim U(10, 100)$). The grey shaded region corresponds to where the total mutational input of both large and midrange alleles is less than 1. The other parameters of the simulation are similar to those in Fig. 4.

Including alleles with midrange effects does not change our qualitative findings about the long-term adaptive response of large effect alleles (*i.e*, with $a^2 \geq 100$). In the parameter ranges we studied, the adaptive contribution of large effect alleles is still maximized near $2NU \approx 1$ and exhibit the same low and high mutational input regimes we discussed in the main for our general case. The qualitative effects on the adaptive contribution are also small, and the behavior of the number of large effect fixations is now simpler in following our low mutational input approximations for a greater range of mutational inputs and in tracking the adaptive contribution (compare Fig. S6 and Fig. S11).

Notably, including midrange effect alleles removes the spike in the number of large effect fixations that we saw for $2NU > 1$ (Fig. S6). This spike is caused by interference: increasing the mutational input increases $D_{qs}$ and decreases $t_{qs}$ (Fig. S8) favoring the fixation of smaller large effect alleles. When alleles with high midrange effects are included, their fixation is favored instead causing a peak in their number of fixations and adaptive contribution (Figs. S11B and E). When the mutational input increases further, the number of fixations with low midrange effects increases alongside their adaptive contribution (Figs. S11C and F). We hypothesize that this peak is partially caused by interference, with $t_{qs}$ being even lower in this range while $D_{qs}$ decreases (Fig. S8), and partially explained by the kind of dynamic described by Hayward and Sella (2022) and summarized in section 4.

We can begin to tease apart the dynamics leading to fixation of midrange effect alleles in the case with one midrange (or large) effect allele alongside a Fisherian background. In Fig. S12, we compare the fixation probability of such a segregating allele in simulations with two alternative approximations. One is the large effect 'double recursion' approximation we introduced in section 5.1, which assumes a stochastic establishment phase followed by a deterministic ascent to frequency ½ (Eq. S16). The other is the non-linear approximation from Hayward and Sella (2022), which assumes an instantaneous deterministic boost in frequency due to the shift followed by the fixation probability with stabilizing selection and genetic drift during the equilibration phase (Eqs. A3-40, A3-48, and A3-16 in Hayward and Sella). We find our approximation to be more accurate for the high midrange and large effect alleles and the Hayward and Sella approximation to be more accurate for small, intermediate and low midrange effect alleles, which accords with our hypothesis.

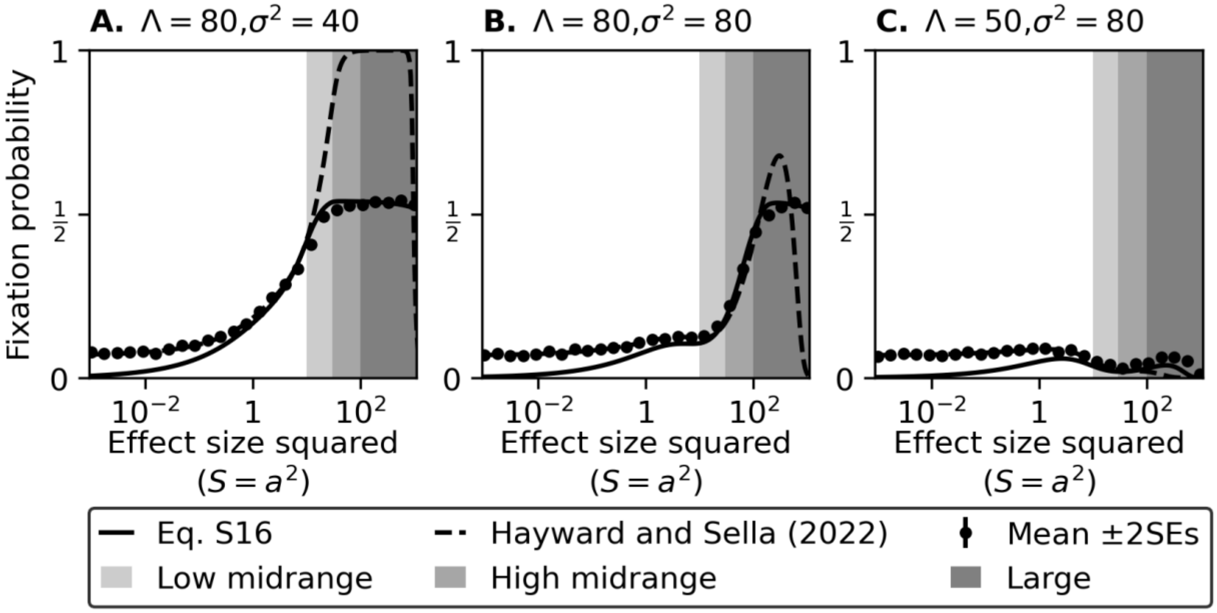

**Figure S12. Probability of fixation for single aligned, segregating allele vs effect size.** We compare simulation results with our approximation based on Eq. S16 using the double recursion approach to find $x_c$ and the non-linear approximation from Hayward and Sella for different shift sizes and background variances. Simulations means and SEs were calculated based on 5000 single-allele simulations. Effect size ranges corresponding to low midrange ($10 \leq a^2 < 30$), high midrange ($30 \leq a^2 < 100$), and large ($a^2 \geq 100$) effect alleles are shaded.

### 7. Phase spaces

#### 7.1. Generating the phase spaces

**Simulation results.** Fig. 6 and related supplementary figures show how the probability of large effect fixations or their adaptive contribution depend on the main parameters of our model or equivalent ones; these figures are projections of the model's phase space. To generate them, we used an even grid along the two axes examined (other than in Fig. 6C and other figures with the same horizontal axis, where we used a finer grid near $p = 0$ and 1), and ran 600 *simplified all-allele* simulations with each combination of parameters in which we recorded the number of large effect fixations and their contribution to adaptation. The contribution to adaptation is given by $2a(1 - x_0)$, where for segregating alleles, $x_0$ is their frequency at the time of the shift, and for new alleles it is $x_0 = 1/(2N)$.

**Smoothing.** Figure S13 shows the simulation results that we used for fig. 6. They are noisy (because of sampling error), so to better visualize them we smoothed the results using a Gaussian filter. Specifically, using $\mathrm{ndimage.gaussian\_filter}(\mathrm{V}, \mathrm{sigma} = [1.5,1])$ from SciPy

(Virtanen et al. 2020), where V is the matrix of simulation results, smooths simulation points based on their ordinal positions rather than their underlying parameter values. We then drew contours based on the smoothed results.

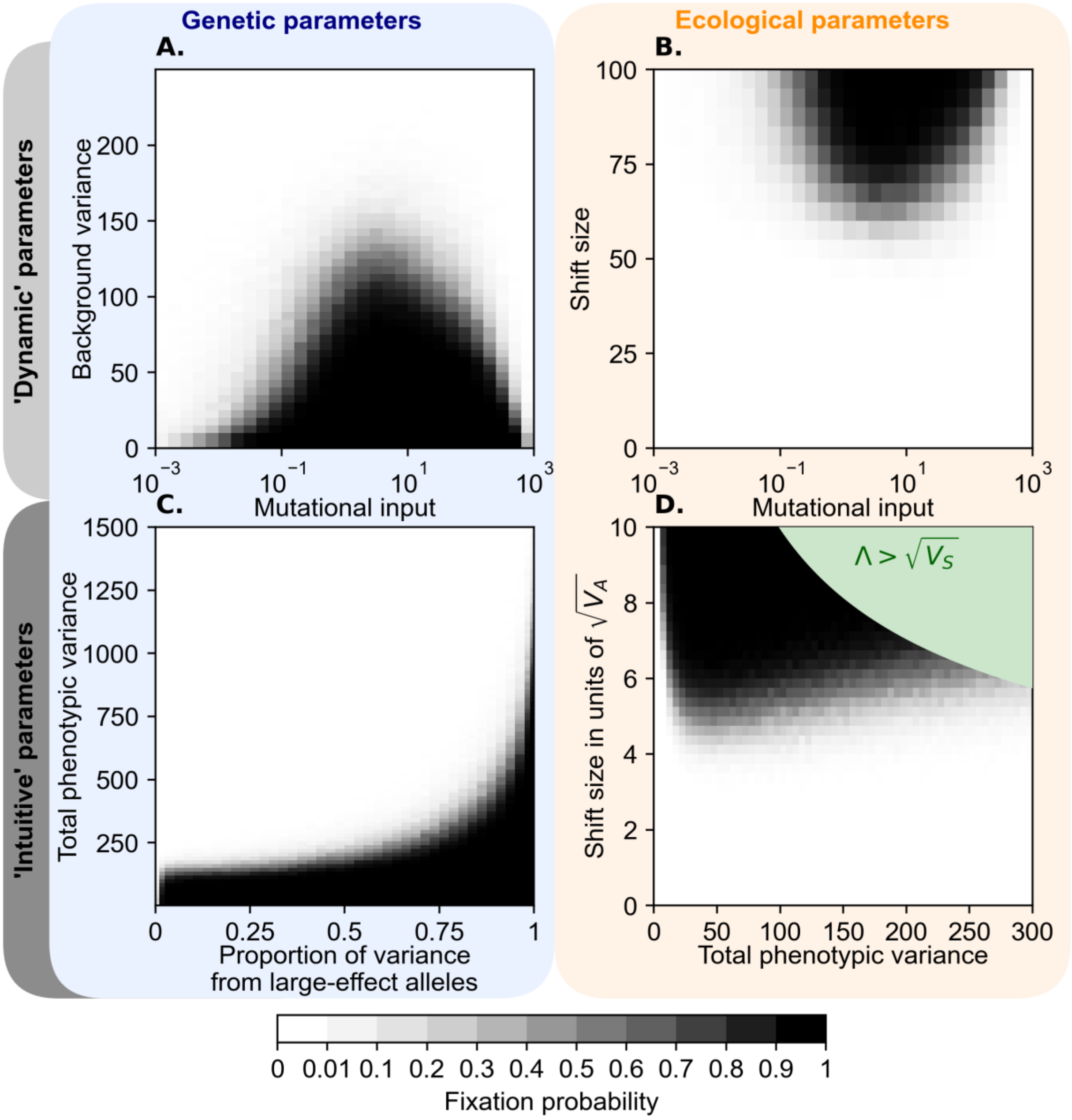

**Figure S13. Phase spaces without smoothing corresponding to Figure 6**. Estimates of the probability that any large effect fixes are shown in squares centered around each grid point.

**Alternative parametrizations.** In Fig. 6C, we replace the genetic parameters we use elsewhere—the background variance ($\sigma^2$) and the mutational input of large effect alleles ($2NU$)—with alternative genetic parameters that are more relatable to observations: the trait's genetic variance at MSDB $\left(V_A(0)\right)$ and the proportional contribution of large effect alleles to this variance $(p)$. The mapping between these parameterizations is one-to-one; specifically, as we already saw the mutational input of large effect alleles and background variance defines the variance at MSDB (Eq. S55) and

$$p = \frac{\int_{10}^{\infty} 2a^2 x(1-x)\tau(x,a)g(a)}{V_A(0)} \approx \frac{8NU}{8NU+\sigma^2}. \tag{S60}$$

#### 7.2. Approximations of contours

We rely on our rough analytic approximation for a single segregating large effect allele to derive rough approximations delineating the parameter values that allow for large effect fixations (Eq. S24). To relate this approximation with our phase spaces, we substitute $\sigma^2 = (1-p) \cdot V_A(0)$ and assume that the allele segregates at frequency $x_0 \approx x_{msdb} = 1/a^2$, which we substitute for $x_c$. In this approximation, fixations can occur if

$$V_A(0) \lesssim \frac{a_{min}(\Lambda - a_{min})}{2\mathrm{Ln}\left(a_{min}^2\right)\cdot(1-p)}, \tag{S61}$$

where $a_{min} = 10$ (corresponding to $a^2 = 100$) and $\Lambda$ is held constant; in terms of shift size relative the phenotypic standard deviation at MSDB, fixations can occur if

$$\Lambda/\sqrt{\mathrm{V_A}(0)} \gtrsim (\mathrm{Ln}(a_{min}^2)/a_{min})(1-p)\sqrt{V_A(0)} + a_{min}/\sqrt{V_A(0)}, \tag{S62}$$

where $p$ is held constant.

In Fig. S14, we compare these approximations with simulation results. Despite ignoring several features of the dynamics, these simple approximations do surprisingly well. For two different shift sizes, we overestimate the variance allowing for fixation near $p = 1$, plausibly because the approximation ignores the effects of interference, and overestimate the variance near $p = 0$, potentially because some large effect alleles start at greater initial frequencies than we assumed (Fig. S14C). For two different proportions of variance from large effect alleles, we overestimate the minimum shift size allowing for fixation at both tiny and large phenotypic variances. We speculate that these deviations also arise from some large effect alleles starting at greater initial frequencies than we assumed.

We use Eq. S62 to calculate the minimal shift size in optimal height that would allow for the fixation of a large effect allele. We assume that (1) $V_A(0) = 3 \times 10^4$, which is an upper bound on the genetic variance contributed by alleles with moderate effect sizes (Simons et al. (2022)) but potentially an underestimate of the total genetic variance (see next section), (2) $p = 0.5$, which is plausibly an overestimate (see next section), and (3) $a_{min} = 10$. This way, we estimate that $\Lambda_{min}/\sqrt{V_A(0)} \approx 40$, where given a standard deviation of ~6.5 cm in standing height (average of within-sex standard deviations, 6.2 cm in females and 6.7 cm in males (Berg et al. 2019; Chen et al. 2022)), suggests that $\Lambda_{min} \approx 260$ cm. As we noted in the main text and given the many assumptions underlying it, this estimate should be taken with a sizable grain of salt.

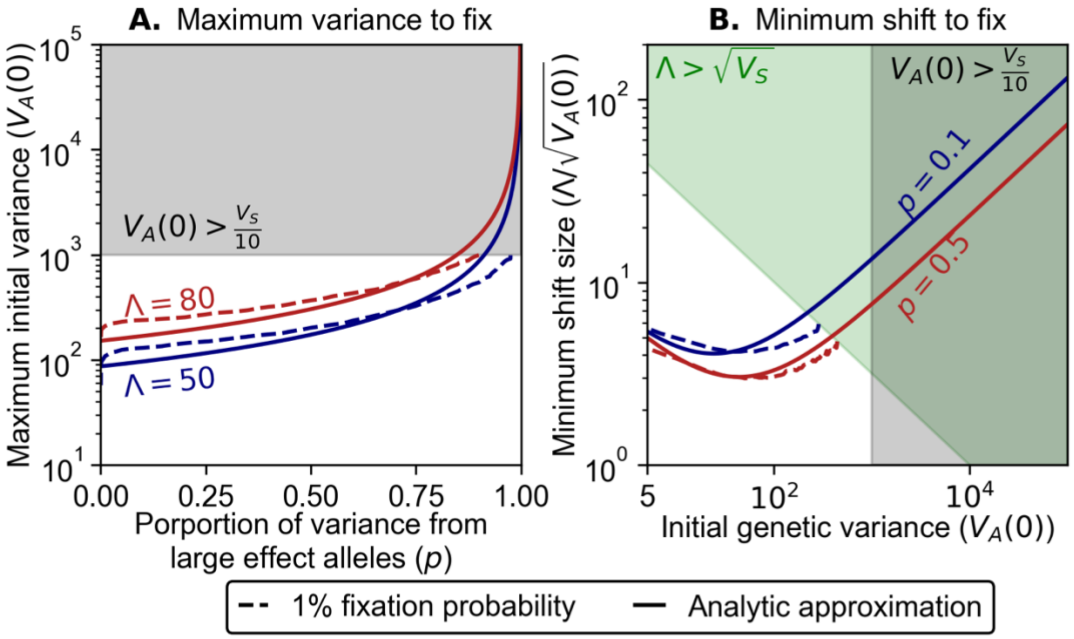

**Figure S14. Simple approximations for parameter ranges allowing for fixation.** We compared our rough approximations (Eqs. S61 and S62) with simulations results from panels C and D in Figs. 6 and Fig. S18. For the simulation results, we used the contours corresponding to a fixation probability of 1%. The green and gray shadings correspond to parameter values that violate our assumptions on the maximal shift size and genetic variance respectively. Simulations results do not extend into parameter regions that violate these assumptions.

### 8. Relating our predictions with empirical findings

#### 8.1. Parameter estimates from human GWAS

In principle, estimates of, and bounds on, the genetic parameters of our model can be inferred from GWAS data. Notably, we can place bounds on the proportion of variance arising from large effect alleles, $p$, based on estimates of the heritability explained by common genetic variation. If we denote the heritability arising from variants with minor allele frequency greater than $q$ by $h_q^2$ then, for example, $p < h_0^2 - h_{0.05}^2 \leq 1 - h_{0.05}^2$, because minor alleles with $a^2 \geq 100$ almost always segregate at frequencies below 5%. The heritability explained by common genetic variation has mostly been estimated in the context of the "missing heritability problem", using lower frequency cutoffs, notably $q = 0.01$. While large effect alleles might sometimes exceed a minor allele frequency of 1%, we plausibly assume that $h^2_{0.01}$ predominantly reflects the contribution of variants with small and intermediate effect sizes. Table S3 shows the bounds on $p$ and the corresponding heritability estimates for a variety of traits. In all cases, $p < 0.95$ and it is typically substantially lower. Moreover, these are plausibly substantial overestimates, because generally $h^2_0 < 1$ and some of the genetic variance arises from rare variants with small and intermediate effects.

**Table S3. Estimates of heritability explained by common genetic variation.**

| Trait(s) | Heritability explained by common variation $h^2_q$ | Frequency cutoff $q$ | Upper bound on $p$ | Reference |
|---|---|---|---|---|
| Height | ~40% | 0.1 | 0.68 | Yang et al. 2015 |
| BMI | ~30% | 0.1 | 0.8 | Yang et al. 2015 |
| Complex diseases | 20%-54% | 0.01 | 0.46-0.8 | Speed et al. 2017 |
| Metabolic traits | 5.6%-37% | 0.01 | 0.63-0.95 | Karjalainen et al. 2024 |

We can also learn about $V_A(0)$ for human traits from the inferences of Simons et al. (2022). Simons et al. estimated three kinds of evolutionary parameters underlying genetic variation in 95 human quantitative traits: the mutational target size $\hat{L}$, the heritability per site in the target $\widehat{h^2/L}$, and the distribution of selection coefficients for new mutations, $\hat{f}(s)$, which they also estimate jointly for all their traits. The 95 quantitative traits were chosen based on having $\geq 100$ approximately independent genome-wide significant hits in the UK Biobank (Bycroft et al. 2018), where this ascertainment criterion biases toward traits with greater values of $L$ and $h^2/L$. The resulting list of the traits includes morphometric, cardiovascular, molecular, blood-related, and ophthalmologic traits and one behavioral trait (Fig. S15).

We can relate $V_A(0)$ of a trait with the estimates of Simons et al. for that trait as follows. We can estimate $V_A(0)$ as

$$\hat{V}_A = \hat{L} \cdot \int E(2x(1-x) \cdot a^2|s)\hat{f}(s)ds, \quad \text{(S63)}$$

where $E(2x(1-x) \cdot a^2|s)$ is the expected contribution to variance per site conditional on it having selection coefficient $s$ and $a$ is the effect size on the trait of an allele at such sites. Under the pleiotropic stabilizing selection model that Simons et al. assume,

$$a^2 = V_S \cdot s/n_e, \quad \text{(S64)}$$

where $n_e$ is the 'effective' degree of pleiotropy of genetic variation in the trait, which is defined as the number of independent traits with the same degree of association with fitness as the focal trait that are required to explain the selection acting on genetic variation affecting the focal trait (supplement section 1.2 in Simons et al. 2018). Given that allele effect size and frequency are independent conditional on the selection coefficient and measuring the trait in units of $\delta$ in which $V_S = 2N$, Eq. S63 becomes

$$\hat{V}_A = \left(\hat{L} \cdot V_S/n_e\right) \cdot \int E(2x(1-x) \cdot s|s)\hat{f}(s)ds = 2Nu \cdot \hat{L} \cdot \widehat{K}/n_e, \quad \text{(S65)}$$

where $u \cdot \widehat{K} = \int E(2x(1-x) \cdot s|s)\hat{f}(s)ds$, $u$ is the mutation rate per gamete per site and $\widehat{K}/n_e$ is the contribution to variance in the trait per unit mutational input.

Figure S15 shows estimates of $n_e \cdot \widehat{V_A}$ for the 95 traits studies by Simons et al. (2020). To generate this figure, we used $u \approx 1.25 \times 10^{-8}$ for the mutation rate per gamete per site (Kong et al. 2012), $N \approx 20{,}000$ for the effective population size (Schiffels and Durbin 2014), and Simons et al. estimates of the mutational target size assuming the SSD model (where the distribution of selection effects is estimated jointly for all 95 traits). The constant $\widehat{K}$ depends on the distribution of selection coefficients and the demographic history of the population. In a constant population with strong selection $\widehat{K} \approx 4$ (see, e.g., Figure 2 in Simons et al. 2018). We used $\widehat{K} = 2.11$, which corresponds to the SSD distribution of selection effects and estimates of demographic history in Simons et al. (2022). Currently, we do not have estimates of $n_e$. On the right horizontal axis of Figure S15 we show how large $n_e$ can be such that we would not expect any large effect fixations in Figure 6C (assuming $p = 0.5$). In the main text, we report the maximal valued of $n_e$ for which we would expect none, or more than half, of the 95 traits to exhibit large effect fixations; we also rely on the estimate of $n_e \cdot \widehat{V_A}$ for height to calculate the shift size at which large effect fixations could occur.

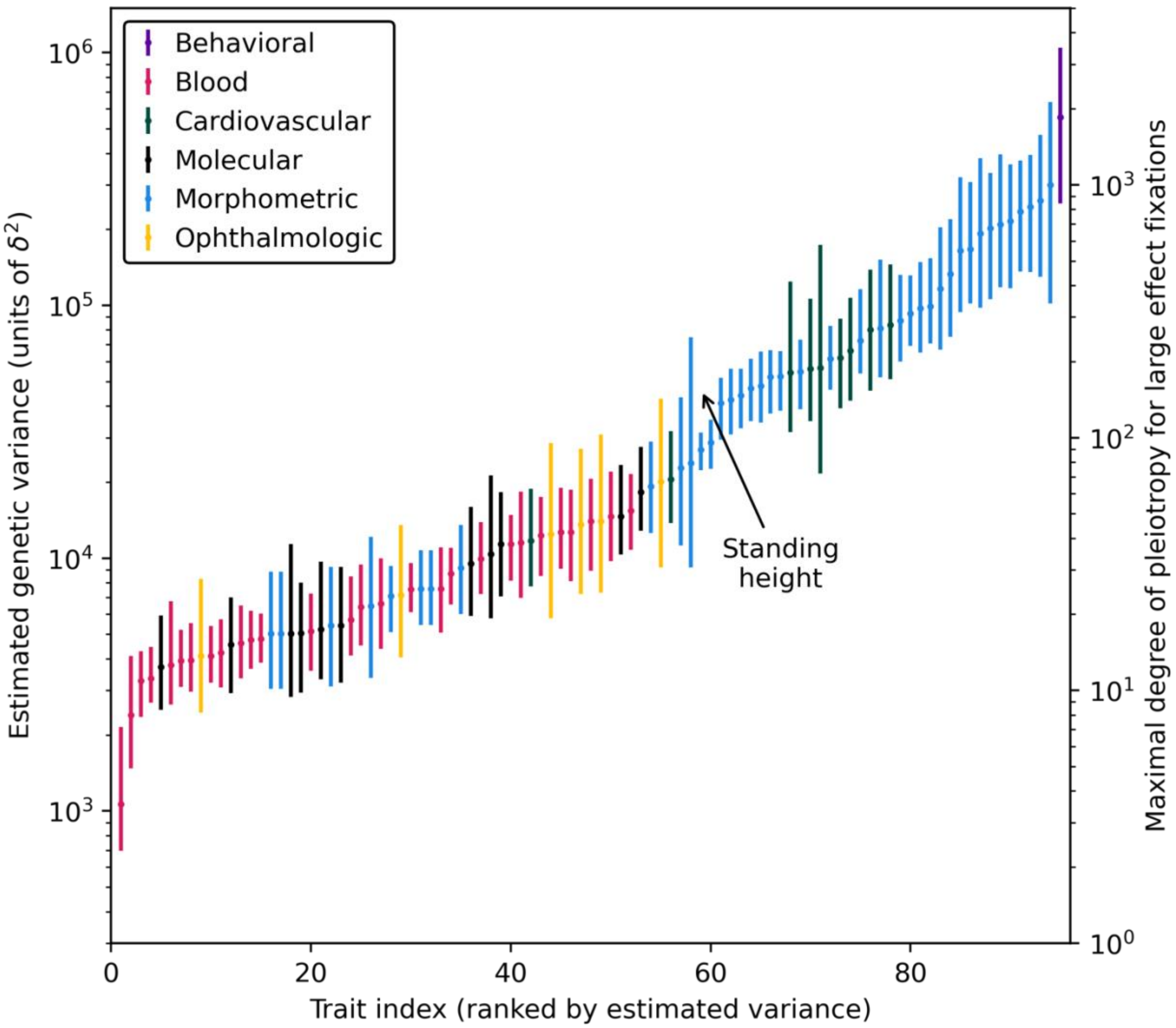

**Figure S15. Estimates of $n_e \cdot \widehat{V_A}$ for the 95 traits from Simons *et al.* (2022)**. Estimates were sorted by value and colored according to six broad categories of traits. See Table S4 for a full list of the traits and estimated values. Error bars correspond to CIs of estimated mutational target sizes from Simons et al. (2022). The degree of pleiotropy required to reduce the genetic variance to 300—the approximate maximal variance that allows for large effect fixations given $\Lambda = 80$ and $p \leq 0.5$ (Fig. 6C)—is shown on the right y-axis.

### 8.2. QTL mapping

We are interested in the overlap between trait ranges in which QTL studies are well-powered to identify large effect adaptive changes and in which we predict they could occur (in, *e.g.*, Fig. 6C). To this end, we consider a QTL study performed with F2 hybrids between two closely related species. We assume that the two species split $T$ generations ago and have had the same evolutionary parameters both before and after they split, *e.g.*, the same population size, distribution of effect sizes and mutational target size for a focal trait. We further assume that one species experienced adaptation to a new optimum of the focal trait, which led to the fixation of a single large effect allele with effect size $a$. We ignore the effects of multiple testing and linkage disequilibrium between markers causal loci on power, as these are difficult to quantify without the specific details of the study.

Under these simplifying assumptions, the power to identify the locus at which the fixation occurred depends only on the proportional contribution of that locus to the phenotypic variance in the F2 hybrids $h_L^2$, the study sample size $\eta$, and a specified probability of type I errors $\alpha$. Accurately calculating power for the QTL analysis depends on the distribution of test statistics under the null (the QTL has no effect) and alternative hypotheses, which are described by a central F-distribution and an F-distribution with non-centrality parameter $n \cdot h_L^2$ respectively (Hu and Xu 2008). For sample sizes of $\eta \gtrsim 50$, power $\mathcal{P}$ is described by a function $\Theta$ that depends approximately only on $\eta \cdot h_L^2$ and $\alpha$, such that

$$\mathcal{P} \approx \Theta(\eta h_L^2|a) \approx \min\big(1, \alpha \cdot \mathrm{Exp}(\eta h_L^2/3)\big), \tag{S66}$$

where by inspection $\mathcal{P}$ increases approximately exponentially with $\eta h_L^2/3$ and is proportional to $\alpha$ but can not exceed 1. The proportional contribution of a locus to phenotypic variance in the F2 hybrids depends both on the locus's contribution to variance $V_L$ and the total phenotypic variance in the hybrid population, $V_T$, where $h_L^2 = V_L/V_T$.

We can derive simple approximations for $V_L$ and $V_T$. Given that the large effect allele fixed in one of the species and is absent from the other, its frequency in the F2 sample would be ½, and its contribution to variance would be

$$V_L = 2a^2 \cdot 1/2 \cdot (1 - 1/2) = a^2/2. \tag{S67}$$

The total variance in the hybrid population consists of four contributions:

$$V_T = V_L + V_{pol} + V_{div} + V_E, \tag{S68}$$

with $V_E$ denoting the environmental contribution to variance, $V_{div}$ denoting the contributions of 'background' small and intermediate effect fixations that occurred in each species, and $V_{pol}$ denoting the contribution of polymorphisms in each species.

We approximate $V_{pol}$ assuming negligible overlap between genetic variation in the two species. In this case, an allele that segregates in one population is expected to segregate at ½ the minor allele frequency in F2 hybrids, such that

$$V_{pol} \approx 2 \cdot 2NU \int_0^\infty \int_0^{\frac{1}{2}} 2a^2 \frac{x}{2}\left(1 - \frac{x}{2}\right) \tau(x|a) g(a) da\, dx. \tag{S69}$$

This expression for $V_{pol}$ approximately equals the genetic variance in each individual species, $V_A$, if we approximate $1 - x/2$ by $1 - x$, which is reasonable given that most minor alleles (and especially those with non-negligible effect sizes) should be rare.

To approximate $V_{div}$, we assume that adaptation negligibly affects the fixation rates of small and intermediate effect alleles, but rather biases fixations in favor of alleles that are aligned with the shift (see, *e.g.*, Fig. S3). The direction of a fixed allele's effect does not change its contribution to variance in F2 hybrids, so we can approximate $V_{div}$ assuming the fixation rate at MSDB. Once again, neglecting overlap in fixations in the two populations and multiple hits we find that

$$V_{div} \approx 2T \cdot 2NU \cdot \int_0^\infty (a^2/2)\pi(a)g(a)da, \quad \text{(S70)}$$

where $\pi(a)$ is the fixation probability of a new allele with effect size $a$ at MSDB (Hayward and Sella (2022)). Under stabilizing selection, minor alleles with non-zero effects are negatively selected, such that large effect alleles practically never fix and the fixation probability of other alleles is less than the neutral fixation probability, $1/2N$.

In Fig. S16, we show how variance explained by the large effect fixation and the sample size required for the QTL study to have 90% power (given $\alpha = 0.95$) depends on the split time between the two species, under three different assumptions about underlying trait parameters. In all cases, we assume that the fixed large effect allele has an effect size of $a^2 = 100$; this is the lowest large effect that we used, so this is a conservative assumption with respect to power. To calculate variance explained and power, we must first calculate $V_T$. We specify $V_A$ from the outset, where per our predictions, we do not expect any large effect fixations for $V_A > 300$ if $p \lesssim 0.5$ in Fig. 6C. Next, we derive $V_E$ assuming the trait has heritability of ½ in the conditions of the assay. Lastly, we require an approximation for $V_{div}$ to approximate the power using Eq. S66. To that end we must specify the distribution of effect sizes. We consider three distributions: (1) in the first example, we assume squared effect sizes for small and intermediate effect alleles are exponentially distributed with mean $a^2=1$, the distribution of large effect sizes used in the main paper, and that 50% of phenotypic variance comes from large effect alleles, (2) in the second example, we assume squared effect sizes are log-uniformly distributed between $a^2=0.01$ and $a^2=1000$, (3) in the third example, we assume the SSD distribution of effect sizes estimated in Simons et al. (2022). Given the distribution of effect sizes, we can calculate the required to mutational input achieve the specified $V_A$ and in turn calculate $V_{div}$, and ultimately the variance explained and power $\mathcal{P}$.

In all the cases we examined, a QTL study with a sample of 300 F2 hybrids should be well powered to identify the large effect locus, so long as the species split time is sufficiently recent. As the split time increases, $V_{div}$ increases linearly due to nearly neutral fixations of small and intermediate effect size alleles, thus reducing the variance explained by the large effect fixation. This reduction is greater when the proportion of new mutations with small and intermediate effects is greater, because we do not expect large effect alleles to fix at MSDB. As a result of this reduction, the sample size must increase linearly with divergence time to maintain a fixed level of power. Additionally, the variance explained by the large effect fixation decreases as $V_A$ increases as such increases in $V_A$ must come from an increase in mutational input for a fixed distribution of effect sizes. In turn, larger mutational inputs increase $V_{pol}$ allow $V_{div}$ to increase faster with divergence time. For $V_A$ small enough that large effect alleles fix frequently ($e.g.$, $V_A \leq 100$), even studies with a sample of 100 or fewer hybrids would be well powered to identify the large effect fixation even in species split for $10N$ generations.

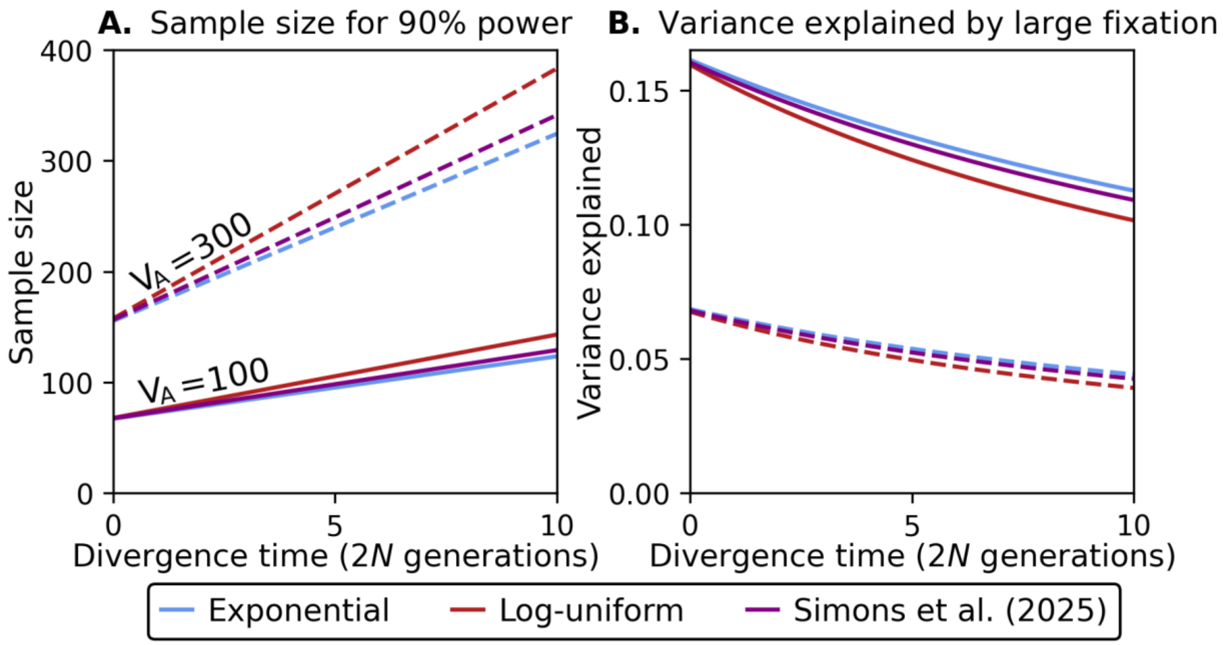

**Figure S16. The power to identify a large effect fixation in a QTL study.** We show (A) the sample size required for 90% power to detect a large effect fixation and (B) the variance explained by the large effect fixation as functions of the split time between the two parent populations for three different distributions of effect sizes and two different genetic variances in each parent population. To calculate these quantities, the phenotypic variance of the hybrid population, $V_T$, was calculated by numerically evaluating S62 and S63 for the given genetic variance in each parent population and distribution of effect size and assuming $V_L = \frac{100}{2}$ , $V_E = \frac{1}{2}V_T$, and $N = 5000$. To calculate power, we assumed a fixed rate of type I errors, $\alpha = 0.05$. Simons et al. (2022) reported a distribution of selection coefficients that we converted to squared effect sizes in units of $\delta^2$ by assuming $a^2 = s \cdot V_S = s \cdot 2N\delta^2$ with $N = 20{,}000$ for humans and truncated the distribution for effect sizes greater than $a^2 = 100$

for which they are underpowered to detect. Additionally, only estimates of the SSD at fixed points were made available for the Simons et al. (2022) SSD distribution. To convert these points to a continuous function, we fit a cubic spline.

### 9. Additional Figures

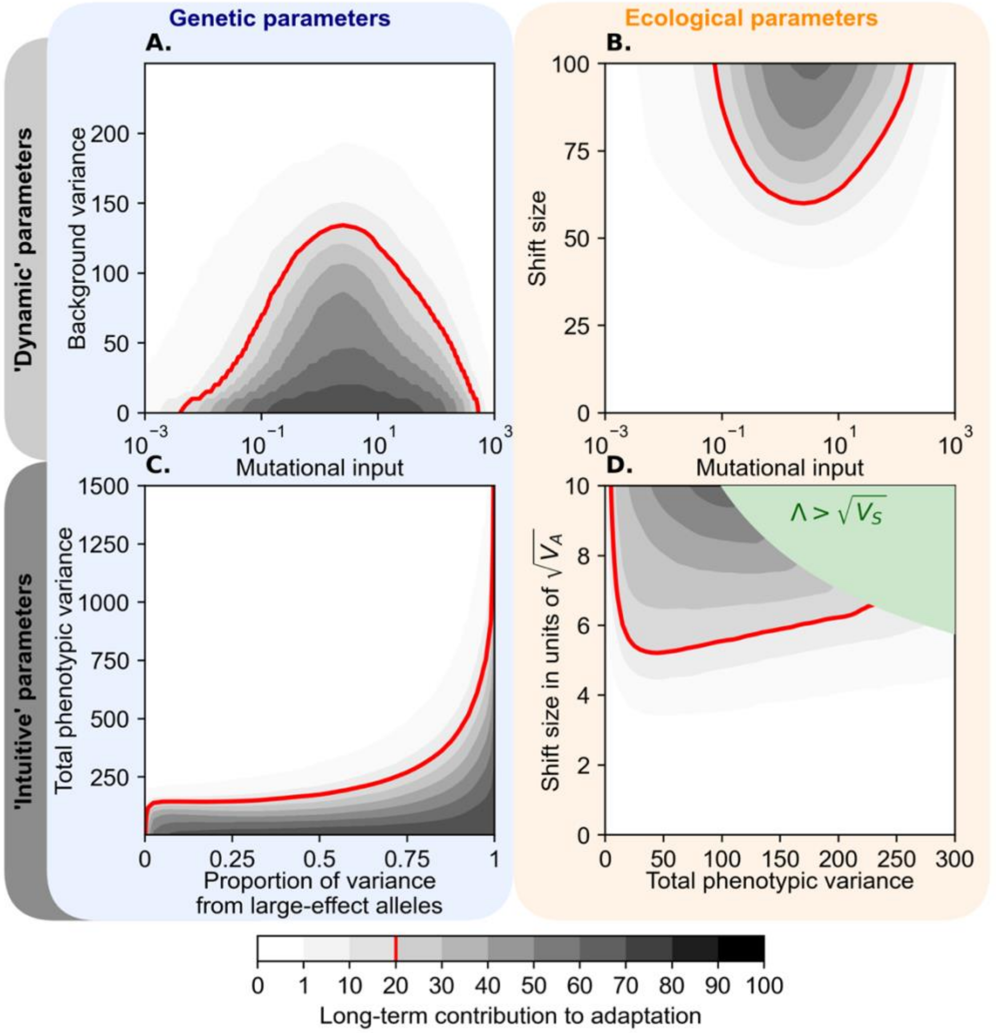

**Figure S17. The adaptive contribution of large effect fixations as a function of the genetic and ecological parameters of traits**. This figure complements Fig. 6, where here we show the adaptive contribution of large effect fixations rather than the probability that such fixations occur. The adaptive contribution is calculated as $\sum_i 2a_i(1 - x_{0,i})$, where the index $i$ denotes a large effect fixation of an allele with effect size $a_i$ and initial frequency $x_{0,i}$, which is the frequency before the shift if the allele was segregating and $1/(2N)$ if it arose after the shift. The adaptive contribution largely mirrors the probability of a large effect fixation, with a secondary effect of the effect sizes and cases in which more than one large effect allele can fix.

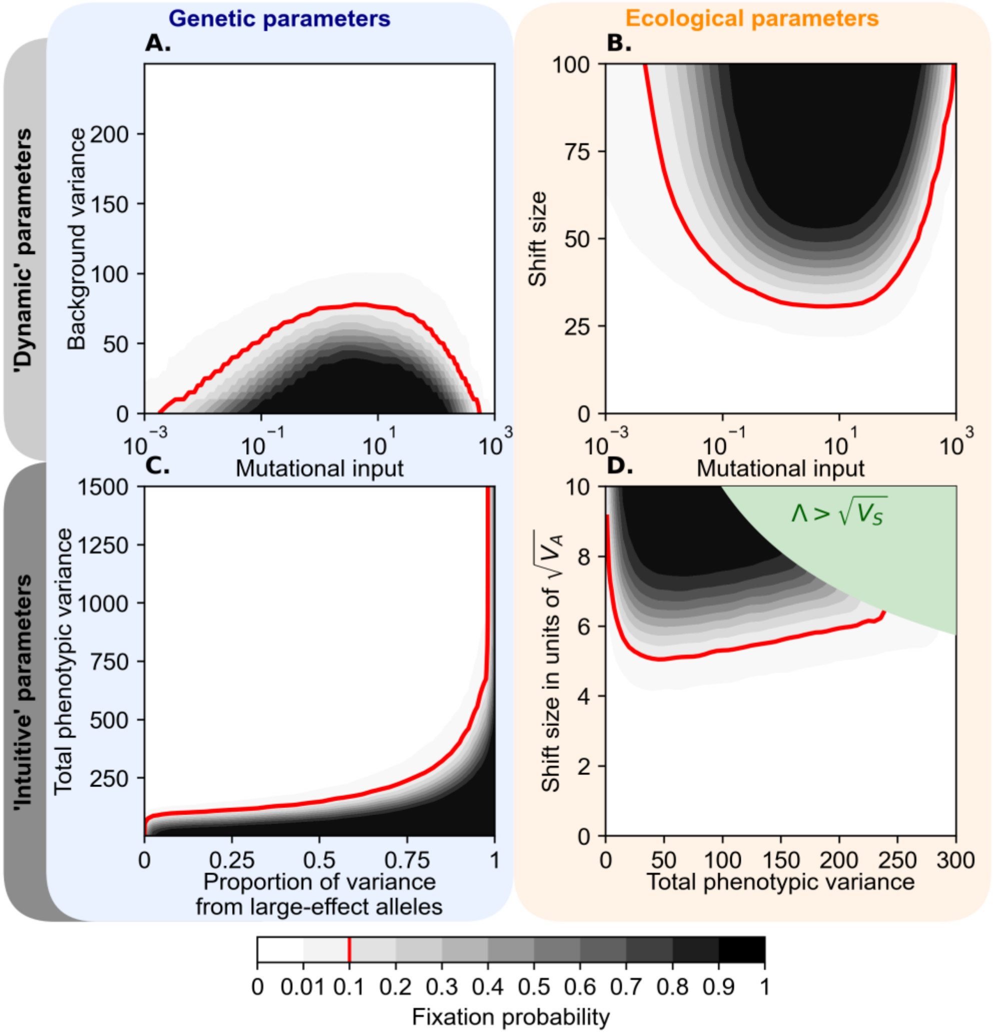

**Figure S18. Alternate choices for fixed parameters in Figure 6.** This figure is similar to Fig. 6, other than having a fixed shift size of $\Lambda = 50$ instead of $80$ in panels (A) and (C), a background variance of $\sigma^2 = 40$ instead of $80$ in panel (B) and a proportion of variance from large effect alleles of $p = 0.1$ instead of $0.5$ in panel (D). A comparison with Fig. 6 indicates that the qualitative behaviors are similar, but large effect fixations are restricted to a smaller region of the parameter space in A, B, and D; large effect alleles fix across a larger region of the parameter space in C, where the background variance is smaller.

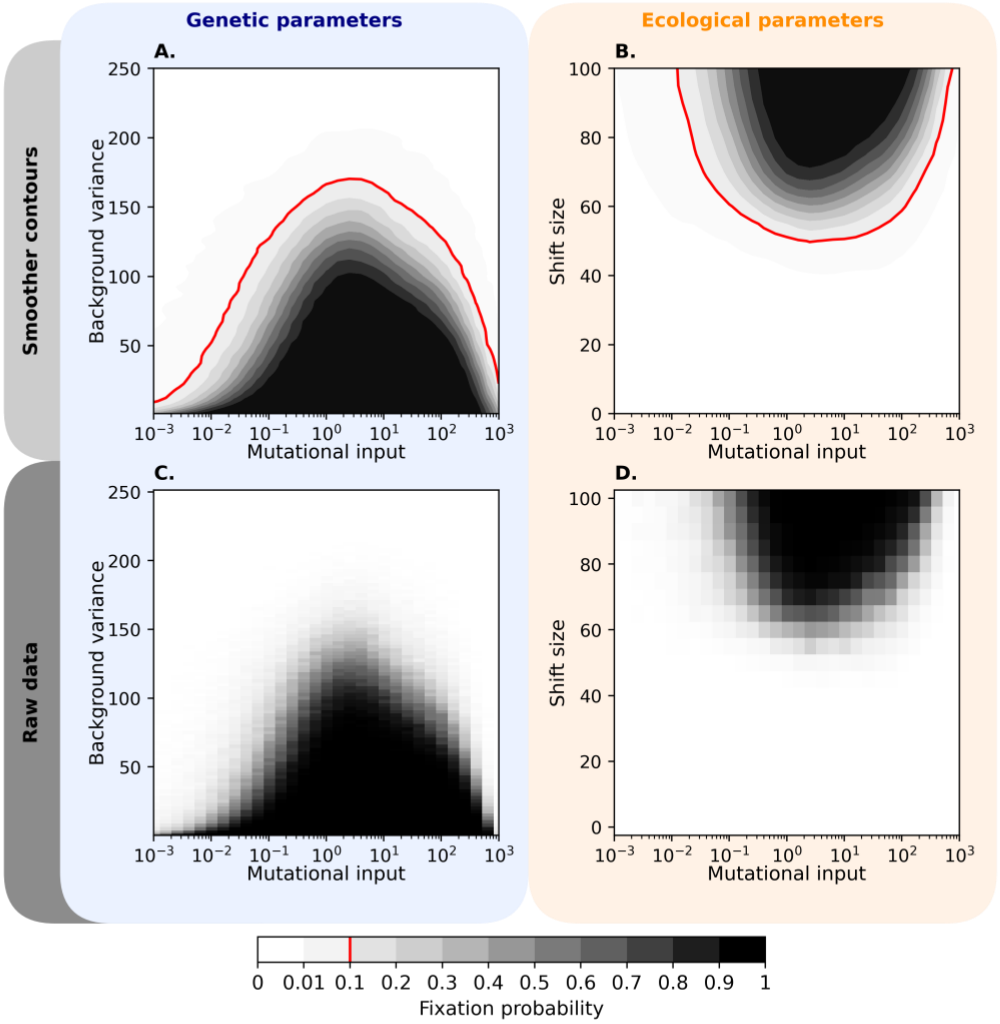

**Figure S19. Alternate distribution of selection effects in Figure 6.** Panels (A) and (B) are similar to the corresponding panels in Fig. 6, where here the mutational distribution of large effect sizes is given by $a^2 \sim U(100, 1000)$ (the alternative distribution in Fig. 1C). Panels (C) and (D) show the same results before smoothing (section 7.1). The results are highly similar to those in Fig. 6, suggesting that they are insensitive to the specific choice of the distribution of large effect sizes (perhaps so long as these are well behaved and/or have similar means).
